# Cell-type-specific nucleotide sharing through gap junctions impacts sensitivity to replication stress

**DOI:** 10.1101/2023.08.15.553366

**Authors:** Benjamin Boumard, Gwenn Le Meur, Marine Stefanutti, Tania Maalouf, Marwa El-Hajj, Reinhard Bauer, Allison J. Bardin

## Abstract

Cell proliferation underlying tissue growth and homeostasis, requires high levels of metabolites such as deoxynucleotides (dNTPs). The dNTP pool is known to be tightly cell-autonomously regulated via *de novo* synthesis and salvage pathways. Here, we reveal that nucleotides can also be provided to cells non-autonomously by surrounding cells within a tissue. Using *Drosophila* epithelial tissues as models, we find that adult intestinal stem cells are highly sensitive to nucleotide depletion whereas wing progenitor cells are not. Wing progenitor cells share nucleotides through gap junction connections, allowing buffering of replication stress induced by nucleotide pool depletion. Adult intestinal stem cells, however, lack gap junctions and cannot receive dNTPs from neighbors. Collectively, our data suggest that gap junction-dependent sharing between cells can contribute to dNTP pool homeostasis *in vivo*. We propose that inherent differences in cellular gap junction permeability can influence sensitivity to fluctuations of intracellular dNTP levels.

**One-Sentence Summary:** The nucleotide pool can be shared between adjacent cells through gap junctions allowing tissue-level buffering of replication stress.

## Main Text

Essential metabolic resources such as amino acids and nucleotides form the building blocks of cell growth and division. How metabolic resources are regulated and distributed in developing and adult tissues is not fully understood. Deoxynucleotide triphosphate (dNTP) production is tightly regulated and defects in dNTP balance can have dire effects, leading to replication stress, DNA damage, and high levels of genetic instability^1–4^. dNTP production can occur via *de novo* synthesis and nucleotide salvage pathways. While *de novo* dNTP synthesis uses amino acids and folate cycle metabolites to produce dNTPs, the salvage pathway relies on precursor deoxynucleosides imported from the extracellular environment or recycled intracellularly from intermediate metabolites^5^. Proliferative tissues use both *de novo* and nucleotide salvage pathways^4–11^.

A critical node of *de novo* synthesis is the enzymatic activity of ribonucleotide reductase (RNR), which catalyzes the rate-limiting step of this pathway. It is exquisitely controlled with its activity peaking in S-phase when dNTPs are required and is further allosterically tuned through binding of dNTPs^12,13^. In addition to this cell-intrinsic control of dNTP levels, external hormones and growth factors activate *de novo* dNTP synthesis, thereby linking local and systemic cues for tissue growth and cell proliferation to dNTP production^14–17^. Thus, a large body of literature has demonstrated that dNTP levels are tightly regulated cell-autonomously with additional important input from extracellular signals leading to "privatized" cell-intrinsic production of dNTPs.

Studies of transformed cells in culture provided evidence of “metabolic cooperation”, allowing for sharing of metabolites which could benefit or harm adjacent cells^18,19^. This cooperative phenomenon occurred between cells in physical proximity and correlated with the presence of gap junctions, leading to the notion that gap junctions underlie this effect^20^. Nevertheless, cell contact-independent transfer of metabolites was also found and other cellular routes including exosomes and nanotubes can also facilitate metabolite transfer between cells^21,22^. Despite these early studies on metabolic sharing of cells in culture, functions *in vivo* of metabolic sharing and its impact on replication stress in developing and adult tissues is not known. Moreover, whether differences exist between tissues in their capacity to share metabolites is not well understood.

Here, we provide *in vivo* evidence of dNTP collective resource sharing at the tissue level depending on exchange of nucleotides from neighboring cells through gap junctions. Our data indicate that a “socialized” mechanism acts within the developing *Drosophila* wing disc allowing tissue-level regulation of nucleotides. This gap junction-dependent exchange provides an important buffering mechanism when dNTP pools are perturbed. However, not all tissues benefit from resource sharing between cells: we find that *Drosophila* adult intestinal stem cells (ISCs) lack gap junctions, have privatized dNTP resources, and are incapable of buffering their dNTP pool when it is depleted. Thus, cells may “privatize” or “socialize” dNTP resources at the tissue-scale, which can impact their ability to respond to changes in dNTP levels. The ability to exchange dNTPs with neighboring cells has important implications for understanding the propensity of cells and tissues to encounter replication stress.

## Results

### Intestinal stem cells are highly sensitive to nucleotide depletion

Previous data suggest a potential role for replication defects in promoting adult stem cell mutagenesis and age-related functional decline^23–27^. We, therefore, further investigated how replication stress affects *Drosophila* adult intestinal stem cells.

Intestinal stem cells (ISCs) replenish terminally differentiated enteroendocrine cells (EEs) and enterocytes (ECs) during homeostasis and in response to tissue damage (Figure 1A)^28,29^. To induce replication stress, we targeted the nucleotide pool through inactivation of *RnrL*, the large subunit of RNR, critical for dNTP synthesis^13^. RnrL was found to be expressed in replication-competent cells in the intestine, with terminally differentiated cells lacking RnrL expression (Figure S1A-B, see also^30^). Knockdown (KD) of *RnrL* in adult ISCs led to high levels of the phosphorylated histone variant and marker of DNA double strand breaks, γH2Av (the *Drosophila* ortholog of γH2Ax) and foci of Replication protein A, associated with replication stress (RpA70-GFP; Figure 1B-E; Figure S1C-E”). Consistent with induction of replication stress, an increased proportion of ISCs in S phase and higher levels of γH2Av in S phase than in G1 or G2 was detected by the FUCCI system, which fluorescently marks distinct cell cycle states^31^ (Figure 1F-I). Feeding adult flies with hydroxyurea (HU), a drug that inhibits RNR, also induced replication stress in ISCs as detected by γH2Av and RpA70-GFP, and blocked stem cell proliferation (Figure S1E-I). Clonal inactivation of *RnrL* in ISCs and their progeny impacted clone growth and induced DNA damage in ISCs as well as other cells in the lineage, which rely on dNTPs for replication in S phase prior to terminal differentiation (Figure 2A-C, Figure S1J-L). Both clonal KD as well as long-term depletion in ISCs of *RnrL* resulted in stem cell loss (Figure 2D-G). Moreover, the ability of stem cells to mount a regenerative response was severely impaired after 2 days of *RnrL* KD (Figure 2H). We conclude that the inactivation of *RnrL* in ISCs has strong effects, causing S phase delays, high levels of DNA damage, stem cell loss, and lineage perturbation. Our data indicate that intestinal stem cells are highly sensitive to replication stress induced by cell-autonomous nucleotide depletion, consistent with known reliance of proliferating cells on the cell intrinsic *de novo* pathway controlling dNTP production^5^.

**Figure 1.**
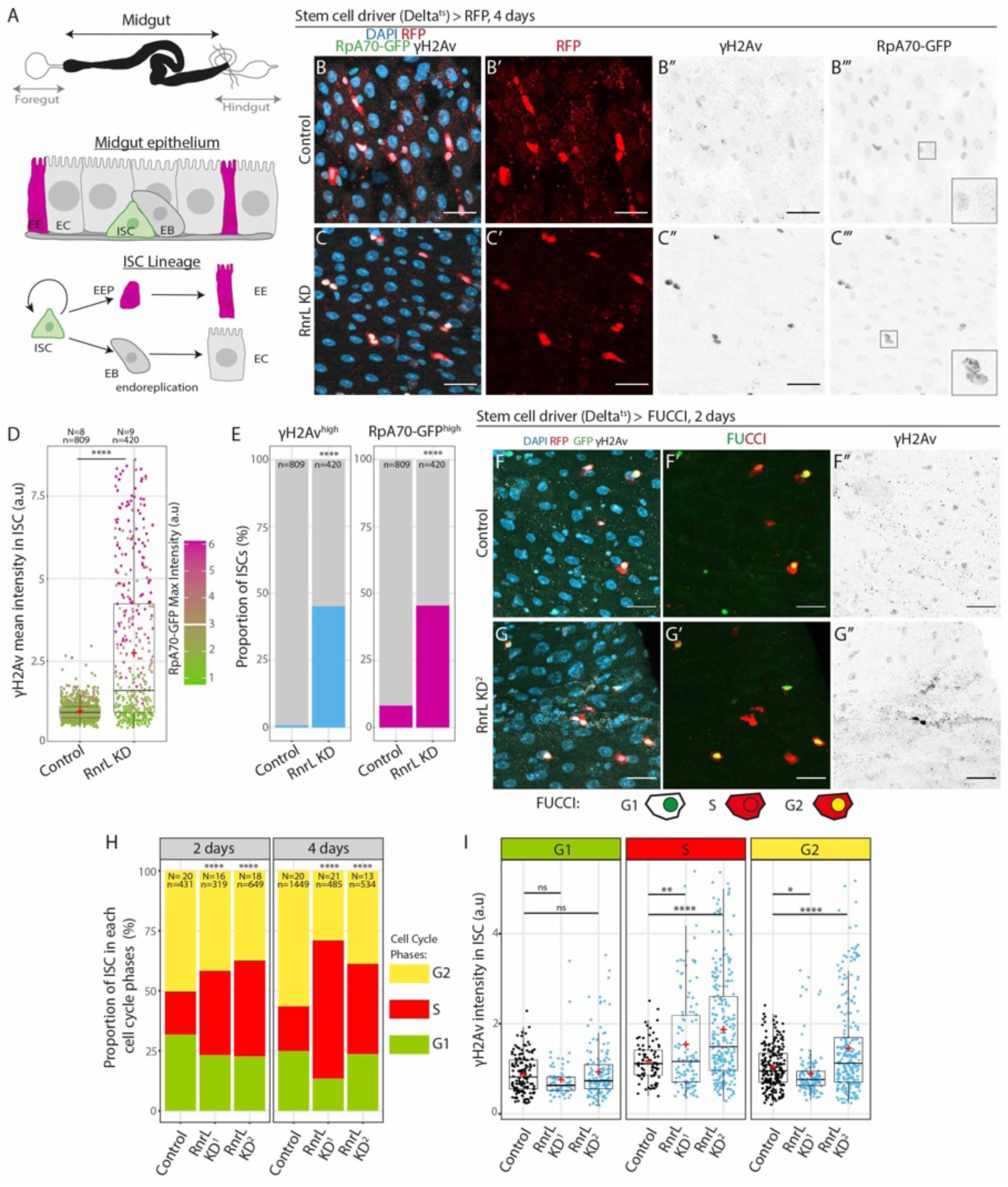
Midgut stem cells are sensitive to replication stress induced by RnrL inhibition. **A.** Representation of the *Drosophila* digestive tract of which the midgut is the main compartment. The midgut epithelium and lineage are composed of Intestinal Stem Cells (ISCs), located basally, that can self-renew and produce Enterocytes (ECs) or Enteroendocrine cells (EEs) through Enteroblasts (EBs) progenitors or Enteroendocrine progenitors (EEPs). **B-C’’’.** *Delta^ts^>RFP, RpA70-GFP* female midguts without or with *RnrL* RNAi for 4 days, DAPI, RFP, γH2Av, GFP are shown. RFP and *RnrL* RNAi expression is specific to stem cells through the *Delta^ts^* driver (*Delta-Gal4*, *tub-Gal80^ts^*). Scale bar: 20µm. **D.** γH2Av mean intensity and RpA70-GFP max intensity in each ISC (RFP+) nuclei in Control or *RnrL* RNAi conditions, normalized to the mean value in the Control. The red crosses represent the mean value. Welch’s t-test, two sided. **E.** Proportion of ISCs (RFP+) with high levels of γH2Av (>2 of mean γH2Av intensity in D) or RpA70-GFP (>3 of max RpA70-GFP intensity in D). Fisher’s test, one-sided. **F-G’’.** Control *Delta^ts^*>FUCCI midguts (F) or expressing *RnrL* RNAi (G) for 2 days. Cells in G1 are marked by nuclear GFP, cells in S phase by cytoplasmic RFP and cells in G2 by both markers. **H.** Proportion of stem cells in each cell cycle phase from *Delta^ts^*>FUCCI with or without *RnrL* RNAi after 2 or 4 days of *RnrL* knock down, two different RNAi lines were used. Chi-squared test. **I.** γH2Av mean intensity in each cell cycle phase after 2 days of *RnrL* RNAi or control in *Delta^ts^*>FUCCI. Normalized to the mean of the Control. Welch’s t-test, two sided, in comparison with control. *: p < 0.05; **: p <0.01; ***: p<0.001; ****: p<0.0001. N = number of guts quantified, n = number of cells quantified, red crosses = mean.

**Figure 2.**
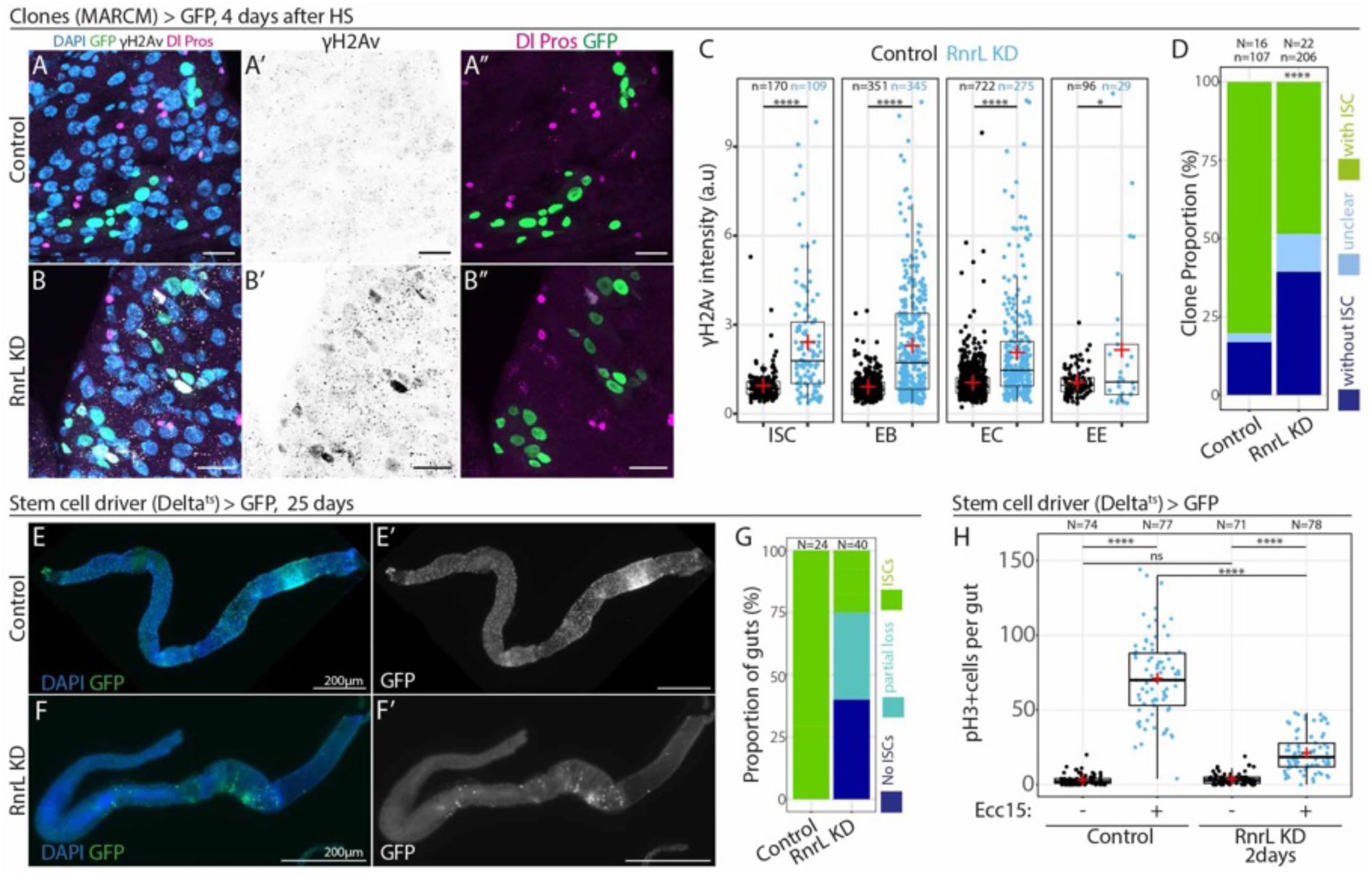
Midgut stem cells are sensitive to replication stress induced by *RnrL* inhibition. **A-B’’**. Control clones (A-A’’’) or *RnrL* RNAi clones (B-B’’’) induced with the mosaic analysis of repressible cell marker system (MARCM>GFP) 4 days after heat-shock (induction); DAPI, GFP, γH2Av, Delta (Dl, membrane vesicles) and Pros (nuclear) are shown. Stem cells are marked with Dl and EEs with Pros. γH2Av indicates DNA damage. Scale bar: 20µm. **C.** γH2Av mean intensity in each cell types in the clones, normalized to Control mean of all cells. Welch’s t-test, two sided, in comparison with control. Clonal depletion of *RnrL* also resulted in high levels of DNA damage, which was especially prominent in ISCs and EBs, but also occurred in ECs and EEs. **D.** Proportion of clones with or without stem cells, 4 days after clone induction in Control and *RnrL* RNAi conditions. Stem cells were detected with Dl. Clones with cells for which the Dl staining was not clearly positive or negative were classified as “unclear”. Fisher’s test, one-sided, comparing the proportion of clones with ISC (green) as part of the total. The proportion of clones without stem cells was increased upon RnrL KD, consistent with ISC loss. **E-F.** *Delta^ts^>GFP* midguts aged for 25 days with or without *RnrL* RNAi. Stem cells were detected with GFP expression. Scale bar: 200µm **G.** Proportion of posterior midguts with normal phenotype or stem cell loss. **H.** PH3+ cells per gut in uninfected conditions or 15 hrs after *Ecc15* bacterial infection of control or *Delta^ts^>RnrL* RNAi flies 2 days after knockdown induction. *: p < 0.05; **: p <0.01; ***: p<0.001; ****: p<0.0001. N = number of guts quantified, n = number of cells quantified, red crosses = mean.

### Wing disc progenitor cells have limited defects upon inactivation of Ribonucleotide reductase

Given our findings of strong defects following nucleotide pool reduction in ISCs, we were surprised to find that wing imaginal disc progenitor cells (Figure 3A), which are highly proliferative during larval stages, behaved differently upon RNR inactivation. In striking contrast to ISCs, a majority of *RnrL* KD clones in the wing disc lacked DNA damage and were indistinguishable from wild-type clones (∼95%, n=506/533), with only ∼5% of clones (+/- 3%, n=27/533) having some detectable γH2Av (Figure 3B-D’). RnrL protein was found to be efficiently depleted by KD (Figure 3E-F’), therefore, why does clonal perturbation of RNR activity have a minimal effect on proliferating wing disc cells?

**Figure 3.**
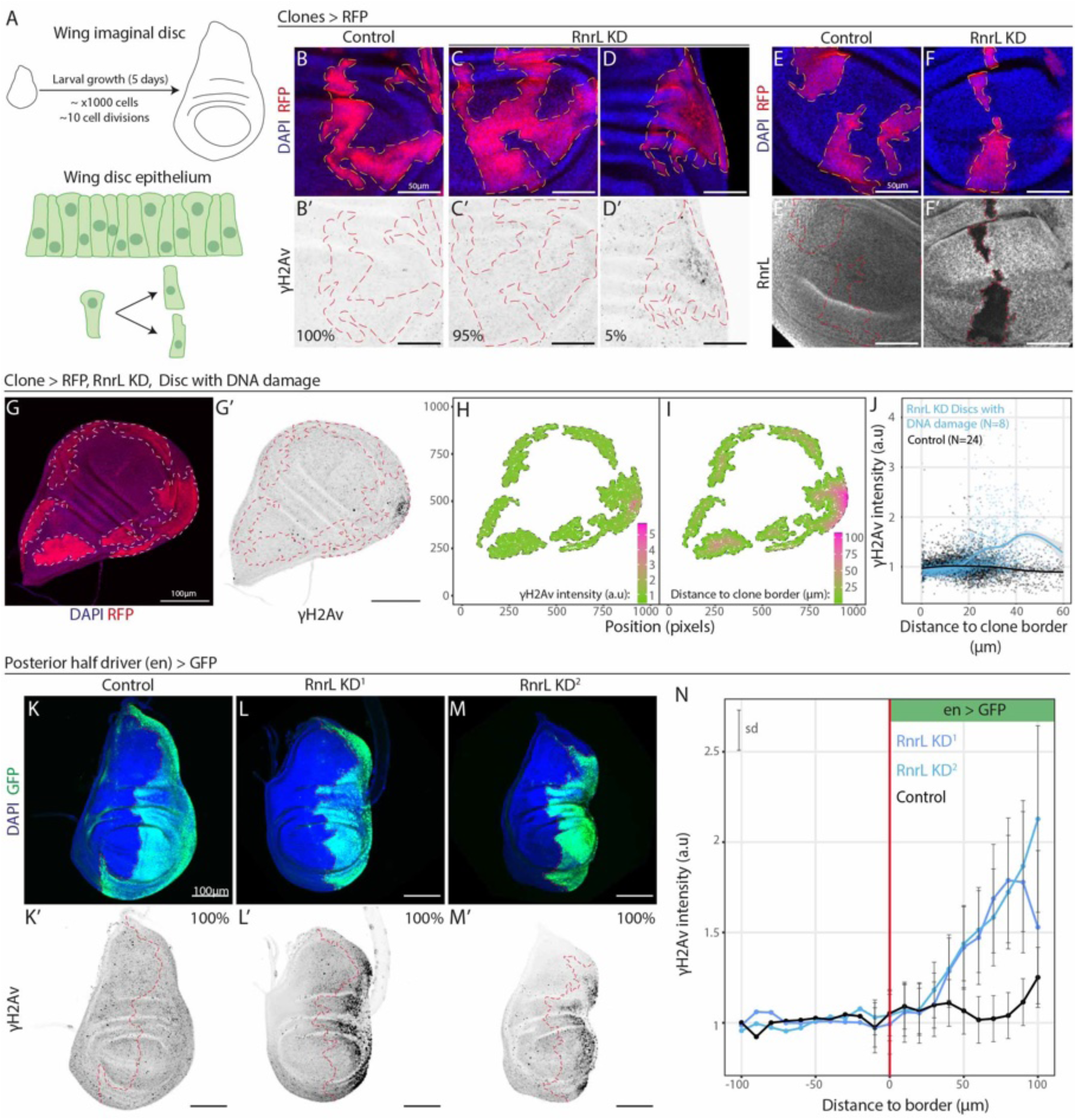
*RnrL* inhibition rarely induces DNA damage in the wing disc. **A.** Representation of the *Drosophila* larval wing disc and wing disc epithelium. **B-D’.** hs-FlipOut>RFP clones in the late third instar larval wing disc, 4 days after induction in first instar larvae, immunostained for RFP and γH2Av, with Control clones (B-B’) and *RnrL* RNAi expressing clones (C-D’). ∼95% of *RnrL* RNAi expressing clones had no detectible DNA damage (C-C’), whereas 5% of *RnrL* RNAi clones displayed a cluster of cells with higher levels of DNA damage at the center of the clone (D-D’). These wing discs and quantifications were obtained from the same experiment presented in Fig 4 D-H’. Scale bar: 50µm. **E-F’.** RnrL protein in the late third instar larval wing disc, 4 days after induction of hs-FlipOut> RFP clones of Control (E-E’) or *RnrL* RNAi (F-F’). Scale bar: 50µm. **G-I’.** Example of quantification method of DNA damage: hs-FlipOut>RFP clones expressing *RnrL* RNAi in the wing disc. (H-I’). In silico mapping of the RFP+ wing disc clones for cells within the clones. The mean γH2Av intensity (H), and closest distance to the non-clonal wild-type tissue outside of the RFP+ clone (I) was measured. **J.** Mean γH2Av intensity per cells in the clones relative to the distance to the wild-type surrounding tissue (clone border; as illustrated in G-I’), in *RnrL* RNAi discs that displayed DNA damage, and Control wild-type wing discs without DNA damage. For each condition, the line is the LOESS curve of local regression (y ∼ x) from the represented dots and the gray area represents a 0.95 confidence interval. The value was normalized to the mean γH2Av intensity in wild-type non-clonal tissue (outside of the clones) for each wing disc. DNA damage was found 20-35µm from the clone border. See individual discs in Figure S3. **K-M’.** Late third instar larval *en>GFP* Control disc (K-K’) or discs expressing two different *RnrL* RNAi constructs (L-M’) stained for DAPI, GFP and γH2Av. The border of the *engrailed* domain (GFP +, posterior) is shown with a red line. Scale bar: 100µm. **N.** γH2Av intensity along the antero-posterior axis and relative to the engrailed domain border in Control and *RnrL* expressing wing discs. For each disc, the γH2Av level was normalized to the level outside the *engrailed* domain and averaged on 10µm intervals. Bars show the standard deviation calculated for each 10µm interval (sd). *: p < 0.05; **: p <0.01; ***: p<0.001; ****: p<0.0001. N = discs.

We considered that activity of the dNTP salvage pathway may produce dNTP precursors, making the *de novo* synthesis pathway non-essential in this context. To test this possibility, we knocked down the *dnk* gene, encoding the sole Deoxyribonucleoside Kinase in *Drosophila*, an essential component of the salvage pathway (Figure S2A-E’). The combined reduction of salvage and *de novo* pathway activity (*dnk* KD + *RnrL* KD) was similar to *RnrL* KD, with a majority of clones having no DNA damage (Figure S2F-L’). In addition, the KD of *dnk* alone in clones did not increase DNA damage, despite efficient depletion of Dnk by RNAi (Figure S2A-B’). Thus, we conclude that the salvage pathway does not have a major role in compensating dNTP precursors in absence of *de novo* synthesis. If the salvage pathway does not compensate for *de novo* nucleotide synthesis, how might wing progenitors undergo replication in absence of RNR activity? One clue came from the ∼5% of *RnrL* KD clones with DNA damage: we noticed that cells with γH2Av were positioned in the center of the clones, 20-35μm from the border with wild-type cells (Figure 3D, D’, G-J; Figure S2M-N). Clones with DNA damage appeared throughout the pouch and wing blade area, with no obvious bias in location (Figure 3A-Q’). Furthermore, when *RnrL* was depleted in the posterior half of the wing disc using the *engrailed-Gal4* driver, high levels of DNA damage were detected on the posterior-most side, ∼40-50 µm from the engrailed domain border with wild-type tissue (Figure 3K-N, Figure S3R-S’). Thus, our data suggest that only when *RnrL* KD cells are far from wild-type cells, do they acquire marks of DNA damage. We therefore explored an alternative hypothesis that a non-autonomous rescue mechanism of nucleotides may arise from neighboring wild-type cells in the tissue.

### Gap junctions can rescue defects of Ribonucleotide reductase depletion

Considering the relationship between the location of cells with DNA damage and the proximity to wild-type tissue, we reasoned that nucleotides might be transferred from wild-type cells to *RnrL* KD cells. Gap junctions, which can allow the passage of small metabolites and ions (<1 kDa), are formed by interactions of transmembrane hemichannels which join between adjacent cells and form pores (Figure 4A)^32^. In *Drosophila*, gap junctions are made by eight Innexin proteins forming homotypic or heterotypic complexes^33,34^. Previous work suggested that the sugar, GDP-L-fucose (∼580 Da), can be transferred between cells of the wing disc in a manner that relies on the gap junction component Inx2^35^. We therefore hypothesized a role for gap junctions in the non-autonomous rescue mechanism in wing disc progenitor cells that can buffer replication stress via transfer of nucleotides (∼500 Da). In the wing disc, the gap junction protein, Inx2 was broadly expressed and reduced upon expression of *Inx2* RNAi (Figure 4B-C’). While *Inx2* RNAi alone did not induce DNA damage (Figure 4D-G’, I-J), co-depletion of *RnrL* and *Inx2* led to high levels of γH2Av in the clonal tissue (91% of clones, n= 243/267, Figure 3D-J). Thus, upon concomitant loss of gap junctions and RNR activity, DNA damage is observed, independent of proximity to wild-type cells. In addition, co-depletion of *RnrL* and *Inx2* resulted in smaller clones than controls, consistent with cell proliferation impairment due to replication problems (Figure 4H, K).

**Figure 4.**
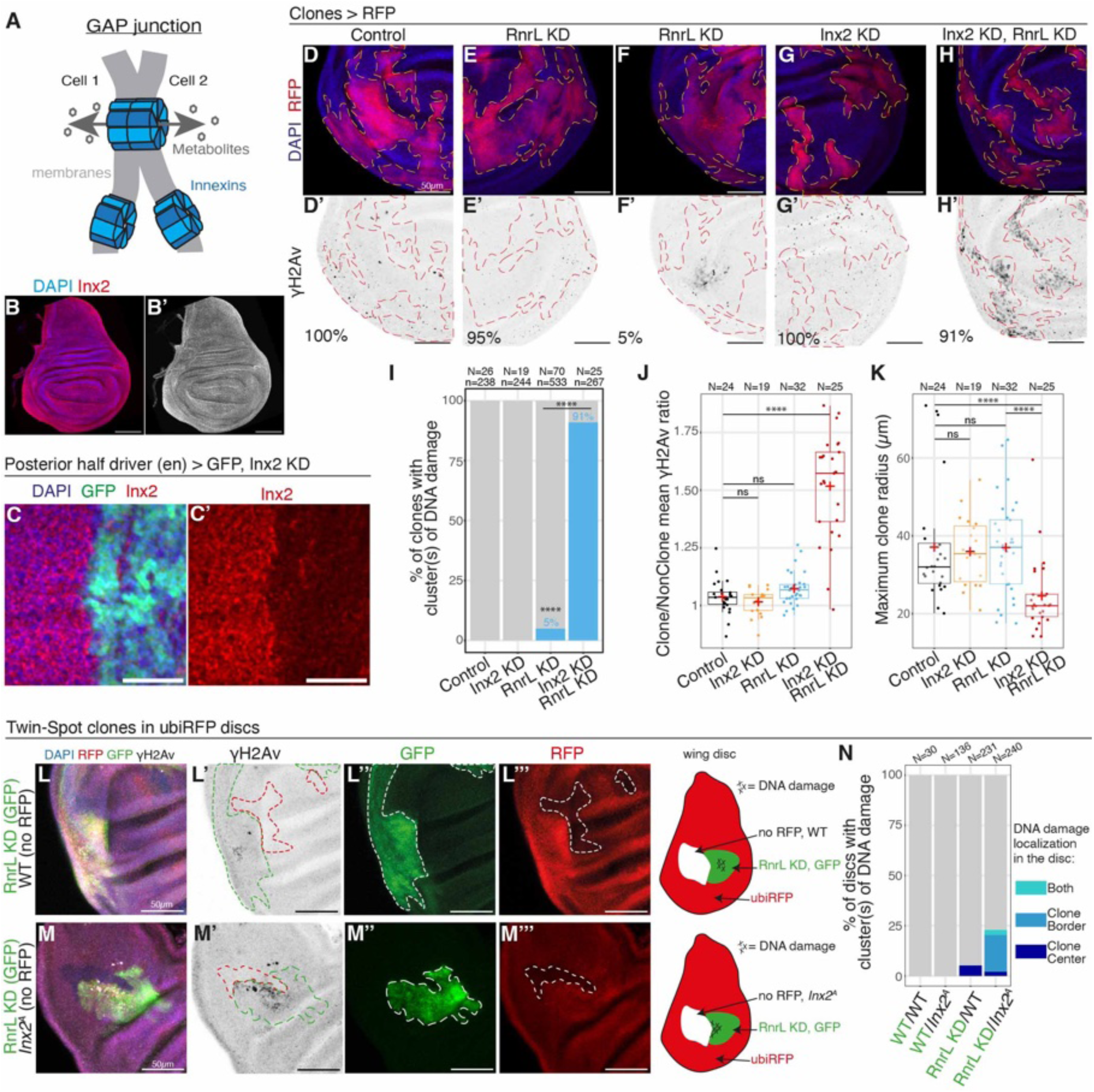
Non-autonomous rescue of DNA damage by gap junctions in the wing disc. **A.** Representation of a *Drosophila* gap junction formed by innexin proteins. **B-B’.** Inx2 protein expression in *Drosophila* late third instar larval wing disc. Scale bar: 100µm. **C-C’.** *en>GFP* wing discs expressing *Inx2* RNAi in the engrailed GFP+ domain, immunostained for Inx2. Close up of the pouch area. Scale bar: 30µm. **D-H’.** Assessment of DNA damage (γH2Av) presence or absence in hs-FlipOut>RFP clones in the late third instar larval wing discs in Control clones (D-D’), *RnrL* RNAi (E-F’), *Inx2* RNAi (G-G’) and *RnrL* with *Inx2* RNAi (H-H’). Scale bar: 50µm. **I.** Proportion of clones with clusters of cells with DNA damage (from D-H’). Fisher’s test, one-sided. **J.** Mean γH2Av intensity ratio between clones (RFP+) and non-clonal wild type tissue for each wing disc analyzed, in hs-FlipOut RFP+ Control, *Inx2* KD, *RnrL* KD or *Inx2* & *RnrL* KD clones. Welch’s t-test, two-sided, in comparison with the control. **K.** Maximum distance (µm) of clone cells to the clone border (WT tissue) in hs-FlipOut RFP+ clones of the wing disc, as a proxy for clone radius/size. Welch’s t-test, two-sided, in comparison with the control. **L-M’’’.** Assessment of DNA damage in Twin spot clones in the wing disc of the indicated genotypes (schematic). As indicated in the schematic, while the tissue expresses RFP ubiquitously, upon clone induction, one clone loses RFP expression and remains wild-type (WT, L-L’’’) or becomes *Inx2^A^* homozygous mutant (M-M’’’), and the twin clone gains Gal4/UAS driven expression of GFP and *RnrL* RNAi. **N.** Percentage of discs with clones displaying clusters of cells with DNA damage (γH2Av), in discs γH2Av was either found at the center of the clone, at the border of the clone near *Inx2^A^* clones or both. See also Figure S4 for additional twin-spot images representing each category. *: p < 0.05; **: p <0.01; ***: p<0.001; ****: p<0.0001. N = discs, n = clones, red crosses = mean.

If nucleotide sharing occurs through gap junctions to neighboring tissue, then blocking gap junctions in cells adjacent to those depleted for nucleotides should induce DNA damage at the border of these populations of cells. To test this, twin-spot clones were produced whereby GFP+ clones expressing *RnrL* RNAi were induced adjacent to clones marked by loss of RFP that were homozygous for *Inx2^A^*, a null allele. Control genotypes did not show DNA damage at the RFP-/GFP+ clone boundary (Figure 4L-L’”, N; Figure S4A-D). Consistent with the above data in Fig. 3C-D’, ∼5.2% of discs with *RnrL* KD GFP+ clones adjacent to RFP-wild-type tissue had γH2Av accumulation in middle of the GFP+ clones (Figure 4N; Figure S4C-D). However, consistent with the transfer of nucleotides from adjacent cells via gap junctions, most *RnrL* KD GFP+ clones with DNA damage had γH2Av accumulation at the border with RFP-*Inx2* mutant tissue (Figure 4M-M”’, N; Figure S4E-H). Together these data support the model that gap junctions between cells buffer levels of nucleotides and replication stress.

Additional evidence for an important role of gap junctions in nucleotide sharing between wing disc cells was found upon knockdown of other enzymes important for dNTP synthesis upstream of RNR (Figure S5A). The concomitant inactivation of gap junctions with KD of either *CTP-synthetase* (*CTPsyn*) or *Phosphoribosyl pyrophosphate synthetase* (*Prps*) led to γH2Av accumulation (Figure 5B-H; Figure S5B-H’, K-L’). Similarly, the inactivation of GMP-synthetase, *bur*, resulted in gap junction-dependent appearance of DNA damage in 20% of discs (Figure 5B, I-J; Figure S5B, I-J’). A similar effect was observed upon KD of *dnk* (Figure S5M-P’). Of note, the knockdown of these nucleotide biosynthesis enzymes alone, did not result in DNA damage, unlike *RnrL* KD, which showed a gradient of DNA damage. This is consistent with RNR activity being a key rate-limiting step in dNTP production. In addition, we note that enzymes with more general roles in ribonucleotide production (Prps RNAi#2, Adenylosuccinate Synthetase – AdSS) also had severe gap junction-dependent growth defects (Figure S5L, L4, Q-R’). DNA damage was not detected in these contexts, likely caused by a primary phenotype of growth restriction due to reduction of the ribonucleotide pool, precluding replication defects that require proliferation. Altogether, these data support gap junction-dependent buffering of nucleotide levels.

**Figure 5.**
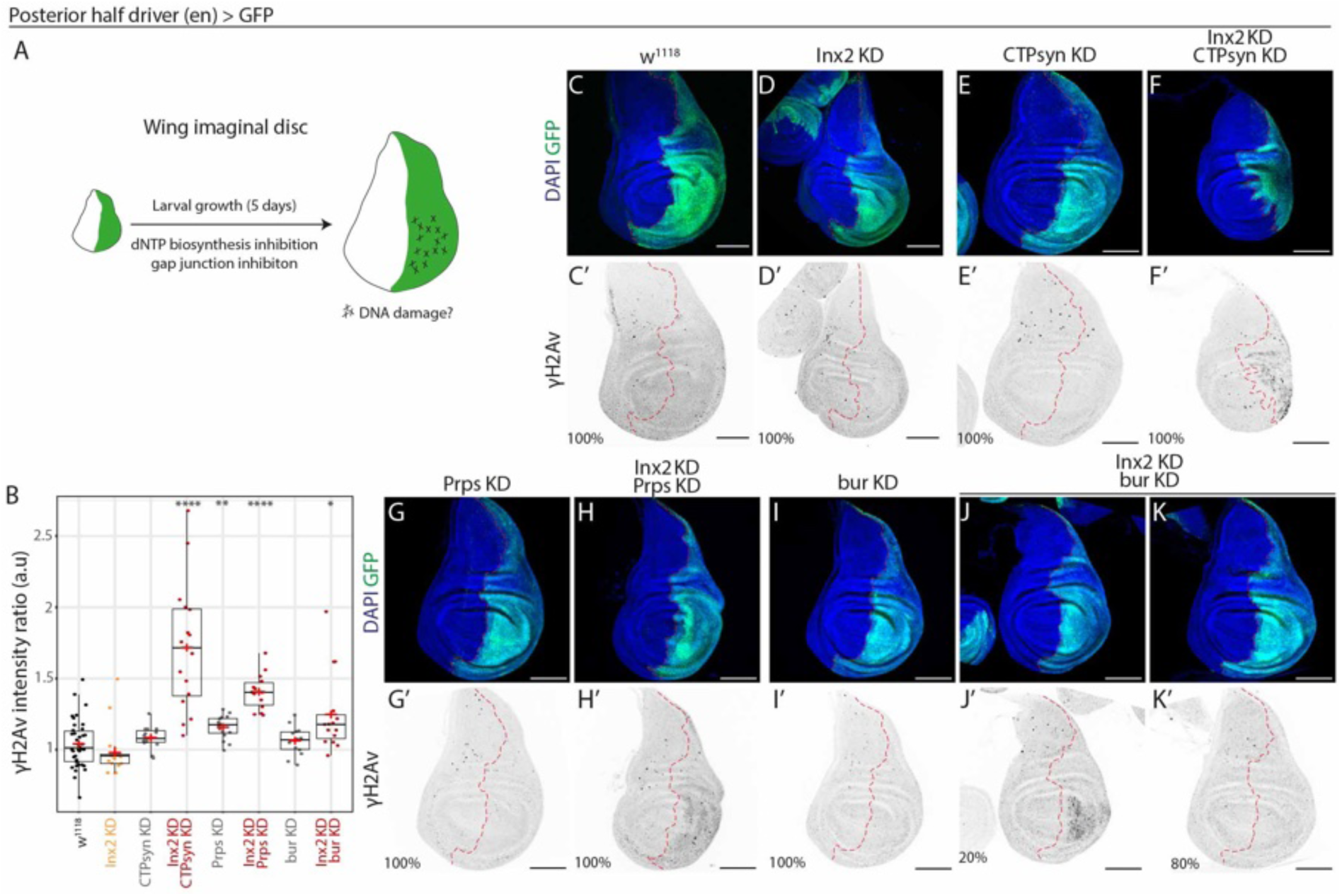
A role for gap junctions in buffering nucleotide between wing disc cells. **A.** Wing disc experimental model. **B.** γH2Av mean intensity in the pouch region of the engrailed domain (posterior, GFP+) relative to the mean intensity in the pouch region of the non-engrailed domain (anterior, GFP-), for the conditions represented in C-K’. Welch’s t-test. **C-K’.** *en>GFP* Control disc (*wildtype*,*C-C’*), disc expressing *Inx2* RNAi (D-D’), *CTP synthetase* RNAi (E-E’), *Prps* RNAi (G-G’) or *bur* RNAi (I-I’), or *Inx2* RNAi in combination with *CTP synthetase* RNAi (F-F’), *Prps* RNAi (H-H’) or *bur* RNAi (J-K’). Late third instar larval wing discs immunostained for GFP marking the domain of expression and γH2Av indicating DNA damage. Scale bar: 100µm *: p < 0.05; **: p <0.01; ***: p<0.001; ****: p<0.0001. Red crosses = mean.

### Intestinal stem cells lack gap junctions

While wing disc epithelial cells can buffer the effects of dNTP pool perturbation through gap junctions, our data above indicated that, in contrast, intestinal stem cells are highly sensitive to depletion of dNTPs upon *RnrL* KD. To understand better the lack of buffering capacity of ISCs, we investigated gap junction protein expression and localization in the adult midgut.

From cell type-specific RNAseq data^36^, we determined that *Inx2*, *Inx3*, and *Inx7* genes are highly expressed in the midgut, with *Inx1* (*ogre*), *Inx4*, *Inx5*, *Inx6*, *Inx8* (*shkB*) expressed at very low levels (Figure 6A-B). In midgut cells, Inx1 and Inx3 appeared to form intracellular puncta and were not enriched at the cell membranes (Figure S6A-F’’), with the exception of a small population of enteroendocrine (EE) cells in the middle midgut that had high levels throughout the cell (R3 region and adjacent regions of R2 and R4, Figure S6G-H’’). Inx8 (shkB) was detectible in some R2 ISCs as puncta, but did not show membrane enrichment (Figure S6I-J’’). Consistent with their very low RNA expression, Inx4, Inx5, and Inx6 proteins were undetectable in the midgut (Figure S6K-M’’). Thus, no cell-to-cell membrane localization, which would be indicative of gap junction formation, was detected with these innexins (Inx1, 3, 4, 5, 6 and 8).

**Figure. 6.**
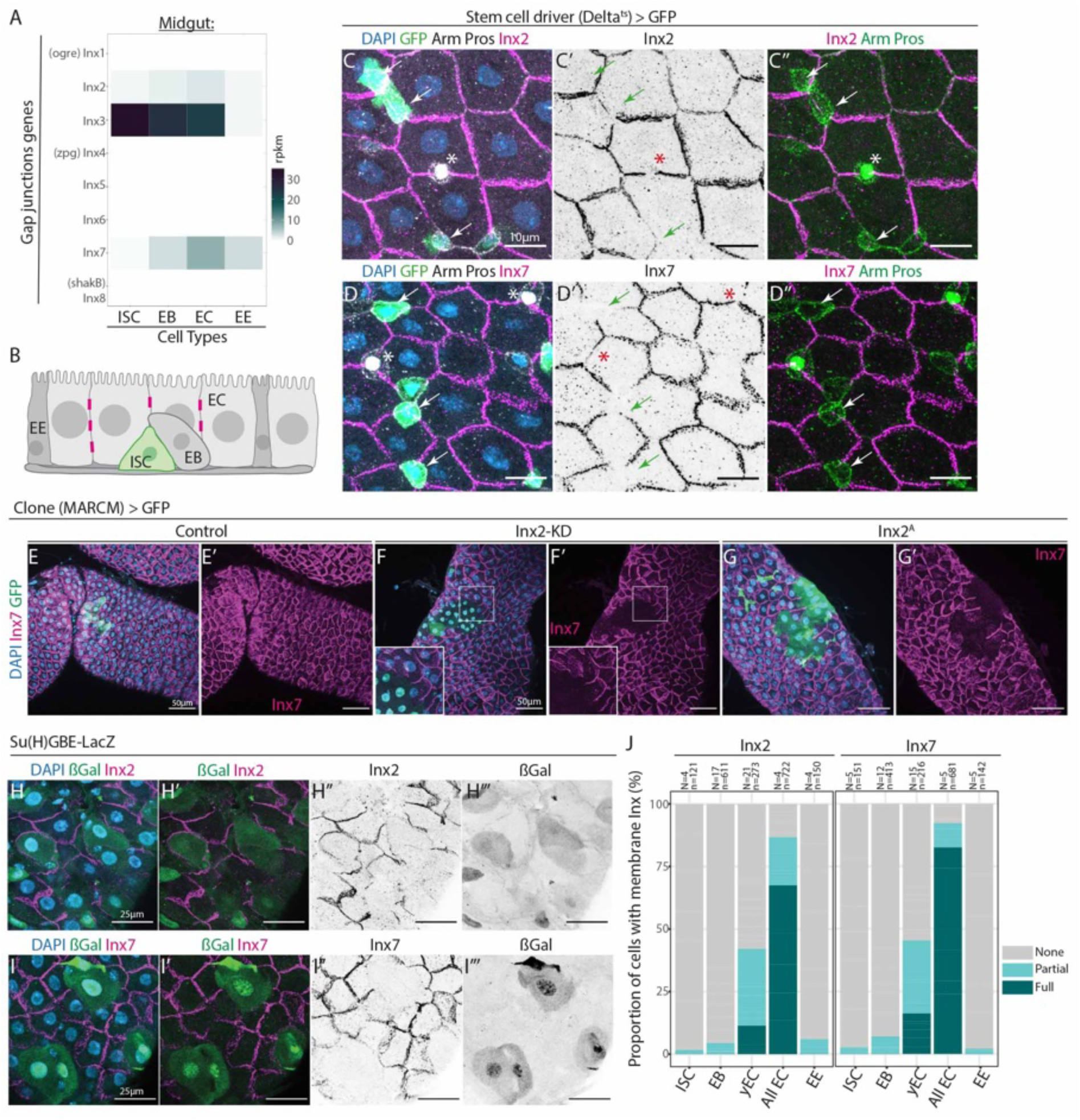
*Drosophila* intestinal stem cells lack gap junctions. **A.** RNA Expression levels of the 8 innexin genes in the intestinal cell types (data from Dutta et al. 2015^36^). *Inx2*, *Inx3*, and *Inx7* genes are significantly expressed in the midgut. **B.** Model of the midgut epithelium, ISCs lack gap junctions (magenta bars). ISC (Intestinal Stem Cell), EB (Enteroblast), EE (Enteroendocrine cell), EC (Enterocyte). **C*-*D’’.** *Delta^ts^>GFP* female midguts marked for DAPI, GFP, Armadillo (membrane), Prospero (nuclear) and Inx2 (C-C’’) or Inx7 (D-D’’). *Delta^ts^* drives GFP expression in the ISCs (white/green arrows), Prospero marks enteroendocrine cells (white/red asterisks), and the ß-catenin Armadillo is enriched at the membrane of diploid cells (ISCs and Enteroblasts). Inx2 and Inx7 were only found at the membrane of large polyploid enterocytes. Membrane location of Inx2 and Inx7 was undetectable in ISCs (arrows), similarly, Inx2 and Inx7 were absent at the membranes of the diploid EBs and Prospero-expressing enteroendocrine cells (asterisks). Scale bar: 10µm. **E-G’.** MARCM>GFP Control clones (E-E’), *Inx2* RNAi clones (F-F’), *Inx2^A^* clones (G-G’) marked for Inx7, DAPI, and GFP. Note the loss of *Inx2* function in clones resulted in absence of Inx7 immunoreactivity in clones and of Inx7 membrane localization in adjacent wildtype cells. Scale bar: 50µm. **H-I’’’.** *Su(H)GBE-LacZ* midguts, Inx2 (H-H’’’) or Inx7 (I-I’’’) as well as ßGal and DAPI are shown. ßGal is expressed in EBs (diploid, small nuclei) and persists in newly made ECs (polyploid, big nuclei). Inx2 and Inx7 were found to be present only between the membranes of mature EC cells and absent from young ECs. Scale bar: 25µm. **J.** Quantification of the proportion of cells with partial or full membrane localization of Inx2 and Inx7 in ISCs (Arm+, ßGal-, small nucleus), EBs (Arm+, ßGal+, small nucleus), young Enterocytes (ßGal+, big nucleus), all Enterocytes (big nucleus) and EEs (Pros+). N = guts, n = cells.

Inx2 and Inx7, in contrast, were strongly enriched at cell membranes of enterocytes (ECs; Figure 6C-D”), consistent with previous data^37,38^. Moreover, the loss of *Inx2* function in clones resulted in absence of Inx7 immunoreactivity in clones and of Inx7 membrane localization in adjacent wildtype cells (Figure 6E-G’). Thus, Inx2 and Inx7 form gap junctions between neighboring ECs and Inx2 is required for Inx7 accumulation in the cell. Gap junctions localized to the basal membranes below septate junctions in ECs (Figure S6N-O’).

Strikingly, membrane location of Inx2 and Inx7 was undetectable in ISCs (Dl>GFP+ cells, arrows Figure 6B-D’’) consistent with low levels of RNA detected in ISC-specific data (Figure 6A, from ^36^). Inx7 and Inx2 were also absent at the membranes of the diploid EBs, enteroendocrine cells (marked by the transcription factor Prospero), and young ECs (Figure 6B-D”, H-J) and showed some regional variation in expression (Figure S7). We conclude that gap junctions in the adult midgut are restricted to mature, differentiated ECs and are absent from ISCs. It is therefore, likely that ISCs rely solely on cell-autonomous production of dNTPs for DNA replication. A lack of ability to receive dNTPs from neighboring cells could explain the vulnerability of ISCs to replication stress induced by nucleotide depletion compared to disc progenitor cells.

### *In situ* visualization of nucleotides supports gap junction-dependent dynamics across the tissue scale

We then wanted to provide direct evidence for gap junction-dependent nucleotide exchange. We took advantage of the fact that 5-ethynyl-2 deoxyuridine (EdU) is a nucleoside analogue that can be tracked in the cell through fluorescent labelling. EdU is imported into the cell through nucleoside transporters and then converted via the salvage pathway (Dnk) into EdU-monophosphate (Figure 7A). Dnk, therefore, is essential to produce an EdU form that can be incorporated into DNA during replication, and the KD of *dnk* should block EdU nuclear staining. However, if gap junctions transport dNTPs between neighboring cells, sharing should occur of the EdU nucleotide produced in the adjacent wild-ype cells with *dnk* depleted cells, thus resulting in EdU nuclear incorporation in the adjacent *dnk* depleted cells (Figure 7A). To test this, we first assessed EdU *ex vivo* incorporation upon KD of *dnk* in the posterior compartment of the wing disc. While control discs (Figure 7B-B’’’) and those expressing *Inx2* RNAi (Figure 7C-C’’’) or a dominant-negative form of Inx2 (RFP-Inx2^DN^, Figure 7F-F’’’) incorporated EdU throughout anterior and posterior compartments, *dnk* KD resulted in a graded appearance of EdU nuclear labelling in the posterior compartment, indicating that EdU-triphosphate was present in these cells despite the lack of *dnk* required to produce it (Figure 7D-D”’, G-G’’’). The EdU nuclear labelling in the posterior compartment of *dnk* KD discs was abolished upon gap junction inhibition (Figure 7E-E”’, H-H’’’), indicating that gap junctions allow the passage between cells of nucleotides. Similarly, clones lacking *dnk* could incorporate EdU in a manner dependent on *inx2* (Figure S8A-B’), indicating transfer via gap junctions from neighboring cells. Thus, tissue-level transfer of nucleotides occurs *in vivo* via gap junctions.

**Figure 7.**
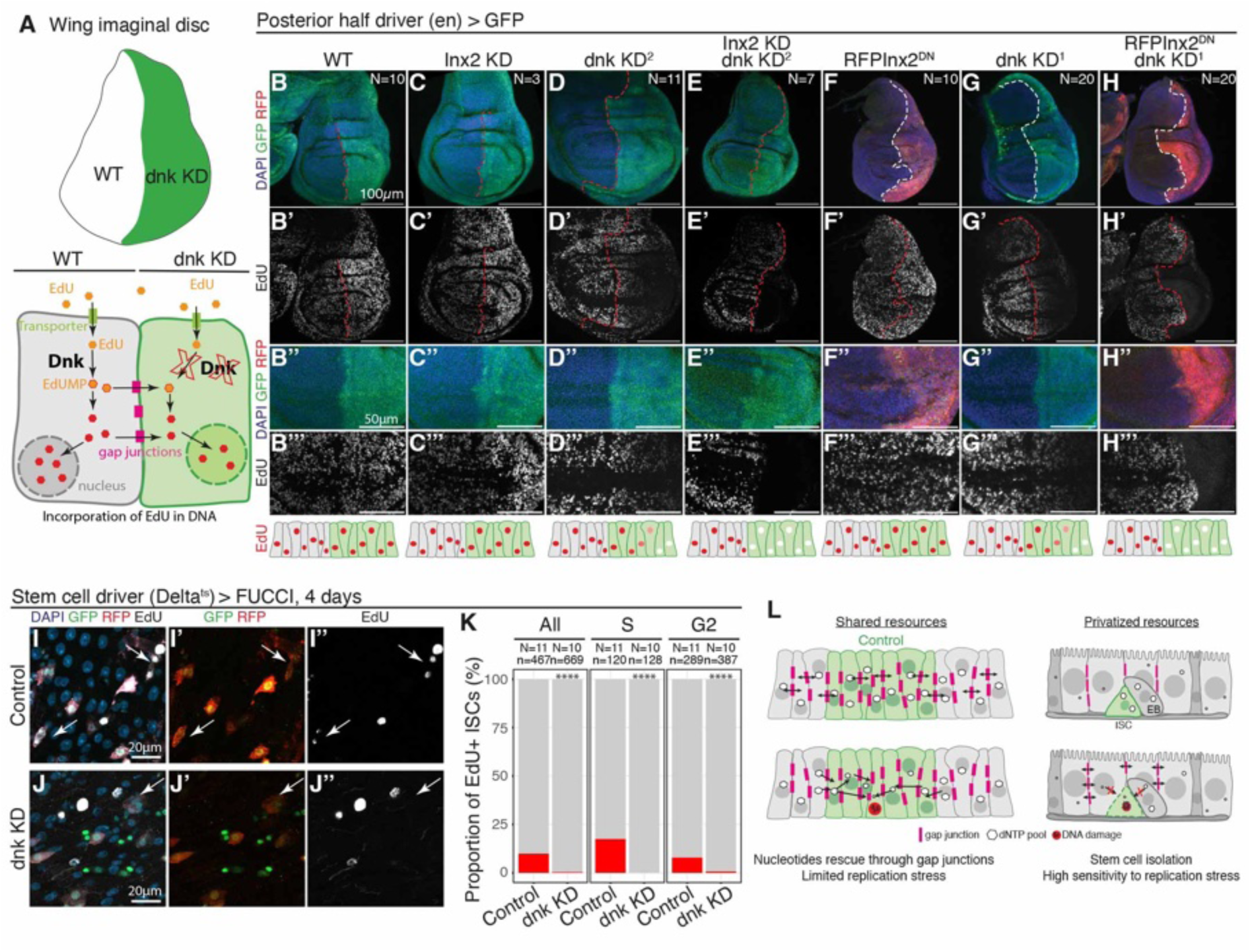
Nucleotide sharing through gap junctions in the wing disc epithelium but not in ISCs. **A.** Model of EdU incorporation in DNA: transport into the cell via transporters of the nucleoside analog, EdU, is followed by phosphorylation by DNK, which is required to produce a nucleotide form capable of incorporation into DNA during S phase. Cells lacking DNK cannot produce the nucleotide form and rely on EdU nucleotides transferred through gap junctions from adjacent wildtype cells for EdU incorporation in the DNA during S phase. **B-H’’’.** *en>GFP* Control disc (*wildtype, B-B’’’*), or discs expressing *Inx2* RNAi (C-C’’’), *dnk* RNAi #2 (D-D’’’), *Inx2* RNAi + *dnk* RNAi #2 (E-E’’’), a dominant negative form of Inx2 (RFPInx2^DN^, F-F’’’), *dnk* RNAi #1 (G-G’’’), or RFPInx2^DN^ + *dnk* RNAi #1 (H-H’’’); labelled for EdU after 30 mins incorporation at the late L3 larval stage. **I-J’’.** *Dl^ts^>FUCCI* Control intestines (I-I”), or those expressing *dnk* RNAi #1 (J-J’’); labelled for EdU after 3 hours incorporation in the adult gut. **K.** Quantification of total, S phase, or G2 EdU+ stem cells. Scale bar: 100µm. N = discs. **L.** Model of nucleotide sharing through gap junctions in different tissues, buffering replication stress in the wing disc progenitor cells but not in midgut ISCs.

ISCs that lack gap junctions would be predicted to be incapable of receiving dNTPs from neighboring cells. To test this, we then compared nuclear EdU incorporation in ISCs expressing the FUCCI cell cycle reporter of a control genotype or KD of *dnk* (to block cell-autonomous phosphorylation of EdU). While 9.9% of control ISCs (Dl-Gal4>FUCCI+ cells) had nuclear EdU incorporation, *dnk* KD ISCs had a very strong reduction (0.4%; Fig. 7I-K). Thus, in contrast to wing disc cells, where gap junctions facilitate phosphorylated EdU transfer, ISCs do not receive phosphorylated EdU from adjacent cells.

Altogether, these findings lead us to propose that the resource sharing of nucleotides through gap junctions provides an important source of dNTPs which can buffer replication stress at the tissue-level (Figure 7L). Moreover, these data suggest that the gap junction connectivity can impact how sensitive cells are to changes in intracellular dNTP and replication stress.

## Discussion

The textbook view of nucleotide metabolism describes their production cell autonomously via the salvage and *de novo* synthesis pathways^6,39^. Our findings reveal an additional abundant source of nucleotides shared from neighboring cells via gap junctions. The ability of gap junction-dependent rescue to compensate for loss of *de novo* synthesis in clones of adjacent cells indicates that it is a robust and potent mechanism acting over large areas of tissue. This favors the idea that nucleotides are being constantly shared between cells connected by gap junctions and thereby comprising a metabolic compartment. This model challenges the current views of nucleotide pools described as being only cell-intrinsically controlled through anabolism and catabolism^5^.

Maintaining the correct concentration of nucleotides is thought to be critical for correctly paced, error-free DNA synthesis^40^. Evidence from yeast and mammalian cells suggests that nucleotides are usually limiting in concentration in cells, necessitating tight regulation throughout the cell cycle, with production peaking in S phase followed by dNTP degradation at the end of mitosis^12^. Early embryos, on the other hand, are pre-stocked with maternal supplies of dNTPs^41,42^. However, slightly later in embryogenesis, dATP concentrations decrease, removing repression of RNR and thereby activating *de novo* dNTP synthesis^41^. This precise control is critical in later embryonic stages as artificially high levels perturb cell cycle kinetics, nuclear movement, and transcriptional output^43–45^. Our findings that nucleotides can be shared between a population of dividing cells therefore raise several important questions pertaining to tissue development and growth: Which cells can donate nucleotides-only those in S phase? Is the concentration of dNTPs equilibrated between the cells of a tissue connected by gap junctions? Does suboptimal dNTP concentrations occur in cells readily exchanging nucleotides and lead to DNA replication or transcription errors? Further investigation of gap junction-dependent sharing of nucleotides within tissues is needed to understand replication at the level of a tissue and contributions of this to development and homeostasis. This could help optimize cancer treatments that act on nucleotide pools perturbation, which may be greatly influenced by this sharing.

Gap junctions have been proposed to subdivide tissues into compartments of metabolite exchange, though this is not well understood *in vivo*^46^. Early studies based on ionic coupling and permeability of injected dyes demonstrated selective connectivity between groups of cells^47,48^. For example, while cells of the early mouse embryos are coupled, boundaries of communication exist between different populations of cells in later stage embryos^49^. Similarly, compartments of restricted dye coupling have been demonstrated in mammalian skin^50–52^. Our data suggest that neighboring cells of the wing disc form a metabolic compartment that can exchange nucleotides. Our findings are consistent with those using dye injection of Fraser and Bryant ^53^, but in contrast to conclusions of previous studies of Weir and Lo^54,55^ suggested that intercellular communication did not cross the anterior-posterior (AP) compartment boundary of the wing disc. We detected innexin-dependent nucleotide exchange within clones throughout the wing disc as well as innexin-dependent EdU passage through the AP compartment boundary of L3 wing discs. Similarly, studies of Ca^2+^ waves using GCaMP signaling found fluctuations moving across the AP boundary of wing pouch in developing and regenerating wing discs^56–58^. Could these differences be due to altered permeability of Luciferase versus nucleotides? Consistent with this possibility, Warner and Lawrence demonstrated that differences exist in cell-to-cell permeability between Luciferase Yellow (less permeable) and a smaller molecule (lead-EDTA, more permeable) over the segmental boundaries in the epidermal cells of milk weed and blowfly larvae^59^. Of note, our data do not exclude potential boundaries between subpopulations of cells within the wing disc. Interestingly, we found evidence that EdU did not transfer from outside the wing pouch to inside this area, suggesting that while globally wing disc cells can exchange dNTPs, local boundaries may nevertheless be present (Figure S8D-G’). This finding was also noted for calcium waves, which were restricted within the folds of the pouch^58^. Thus, subtle differences in size or other molecular features as well as altered gap junction expression or tissue constraints could impact the limits of metabolic boundary compartments. Furthermore, it is also possible that in some cell types, larger molecules such as GFP (Mw=25kDa) may pass through gap junction like-pores as recently demonstrated in polyploid cells of the *Drosophila* hindgut^32,60^.

Additional studies have begun to unravel aspects of gap junction-mediated metabolic exchange^61^, though these are primarily focused on calcium due to its ease of detection with sensors such GCaMP^62^. The innexin-dependent propagation of calcium waves between cells is detectable in numerous tissues including glia, neurons, epidermal cells, imaginal discs progenitors, enterocytes, and the lymph gland in *Drosophila*^37,38,56–58,63–67^. Of note, stem cells are often metabolically linked to neighboring cells with gap junctions, including mammalian skin stem cells^68^ and *Drosophila* germline stem cells^69^. It is somewhat surprising then that *Drosophila* ISCs do not have gap junction connections to neighboring cells. This may be related to the need to isolate stem cells in gut epithelia from their adjacent differentiated cells that have diverse metabolic functions and likely have very different concentrations of metabolites. Overall, our findings underline that tissue-level metabolite exchange has important implications for understanding homeostasis of developing and growing of tissues. Given their increased number of gap junction protein-coding genes (22 connexins in humans vs 8 innexin genes in *Drosophila*) and possible combinations of gap junction assembly, it is likely that mammalian tissues have additional levels of regulation on metabolite exchange, consistent with a recent preprint^70^. Our study, therefore, provides a starting point for future investigation to better understand the molecular features and tissue barriers controlling metabolite flows within tissues and the impact this has on development and homeostasis.

## Acknowledgments

We would like to thank the members of the Bardin team, P.-A. Defossez, Y. Bellaïche, P. Leopold, R. Basto, K. Siudeja, and C. Desplan for feedback on the manuscript. We thank G. Ammer, A. Arnaoutov, G. Tanentzapf, A. Lalouette, P. Spéder, J-R. Martin, P. Leopold, Y. Bellaïche, and M. Dasso for antibodies and *Drosophila* lines as well as the Bloomington and Vienna stock centers for fly lines and Flybase for invaluable information. We thank Prof. M. Hoch for his support of innexin research at the Limes institute. Microscopy training and access has been provided by The Cell and Tissue Imaging Facility/UMR3215 (PICT-IBiSA) of the Institut Curie, members of the French National Research Infrastructure France-BioImaging, ANR10-INBS-04.

## Funding

Work in the Bardin lab is supported by Fondation pour la Recherche Médicale (A.J.B., EQ202003010251), the program “Investissements d’Avenir” launched by the French Government and implemented by ANR, ANR SoMuSeq-STEM (A.J.B., ANR-16-CE13-0012-01), ANR ChronoDamage (A.J.B., ANR-20-CE13-0013_01), ANR Enviro-Print (A.J.B., ANR-23CE13-0024-01), ANR Infinitesimal (A.J.B., ANR 23CE15003802), Labex DEEP (ANR11-LBX-0044), IDEX PSL (ANR-10-

IDEX-0001-02 PSL). Salary support for A.J.B. was from the CNRS and B.B from ENS de Lyon, FRM (FDT202001010957). R.B was supported by the SFB 645.

## Author Contributions

Conceptualization: BB, AJB.

Methodology: BB, AJB.

Reagent contribution: RB.

Investigation: BB, GLM, TM, ME-H, MS.

Visualization: BB, MS, GLM, AJB.

Funding acquisition: BB, AJB.

Project administration: AJB.

Supervision: BB, AJB.

Writing – original draft: BB, AJB.

Writing – review & editing: BB, RB, AJB.

### Competing interests

Authors declare that they have no competing interests.

### Data and materials availability

All data are available in the main text or the supplementary materials, and all reagents can be made available from the corresponding author upon request.

## MATERIALS AND METHODS

### Fly Stocks

The experiments presented in this manuscript used the following fly lines.

From the Bloomington stock center: *UAS-2XGFP* (BL6874), *UAS-nlsRFP* (BL30556)*, UAS-RnrL RNAi* #1 (BL51418), *UAS-RnrL RNA*i #2 (BL44022), CTPsyn RNAi #1 (BL53378), UAS-CTPsyn RNAi #2 (BL31752), UAS-UAS-CTPsyn RNAi #3 (BL31924), UAS-Prps RNAi #1 (BL60086), UAS-Prps RNAi #2 (BL35219), UAS-dnk RNAi #3 (BL65886), UAS-dnk RNAi #4 (BL35219), UAS-bur RNAi #1 (BL60432), UAS-bur RNAi #2 (BL31055), *Inx2^A^ FRT19A* (BL54481), UAS-AdSS RNAi (BL33993).

From the Vienna Drosophila Resource Center: UAS-dnk RNAi #1 (v39137), UAS-dnk RNAi #2 (v103385).

The following stocks were gifts: *w^11^*^18^ (M. McVey); *RpA70-GFP* ^31^ (E. Wieschaus); FUCCI ^31^(B. Edgar); *En-Gal4* (Y. Bellaiche, P. Leopold); *nub-Gal4* ^71^ (P. Leopold) ;*UAS-Inx2RNAi and UAS-RFPInx2^DN^* ^64^(P. Spéder); Su(H)GBE-LacZ ^72^ (S. Bray); *MARCM19A* and *FRT19A* (A.Gould); hs-flp; ; *tub-FRT-GAL80-FRT-GAL4*, *UAS-mRFP/TM6b, Tb, Hu* (originally gift of E. Martin-Blanco to Y. Bellaiche; ^73^); *UAS-rnrL-P2A-rnrS*^30^ (M. Dasso).

The *Delta*^ts^ line was generated from the *DeltaGal4/TM6TbHu* (S. Hou). It was used to generate: *UAS-2xGFP; Delta^ts^/ TM6TbHu*; *UAS-RFP;Delta^ts^/ TM6TbHu*; *FUCCI; Delta^ts^/TM6TbHu*.

### The flies used for the TwinSpot experiment were generated using stocks from the Bloomington stock center: BL42726, BL31406, BL54481, BL51418

The table S1 summarizes all the stocks used, and the table S2 indicates the genotypes used and represented in each figure.

Unless indicated the flies were kept at 25°C on standard medium with yeast. For the temperature sensitive experiments, flies were crossed at 18°C and the progeny kept at 18°C until 2-3 days old adult were collected and transferred at 29°C for the time of the experiment (2, 4, 8 or 25 days) before dissection, with fresh medium provided every 2-3 days.

### HU feeding

For the hydroxyurea (HU) feeding experiments, 3-4 days old flies were placed on food supplemented with sterile distilled water (Control) or 10 mg/mL of HU diluted in sterile distilled water.

### Clone Induction

<u>MARCM in the gut:</u> For the clone induction in the gut, adult flies 2-3 days after eclosion were heat-shocked at 36.5°C for 30 mins and then kept at 25°C on yeasted food for the duration of the experiment. <u>FlipOut and TwinSpots in wing disc:</u>

For the clone induction in the larval wing discs, adult flies were left on yeasted food at 25°C to lay eggs for 24 hours. Progeny were heat-shocked at 37°C for 1h, 24 hours after the end of the egg laying period, and then kept at 25°C for the duration of the experiment. The wing discs were dissected 90-96 hours later, at late L3 stage (wandering larvae).

### Immunostaining

#### Guts

The tissues were fixed and stained following a previously published protocol ^74^. Briefly, the flies were dissected in Dulbecco’s PBS (Sigma-Aldrich D1408) and the tissues were fixed at room temperature in 4% paraformaldehyde for 2 hrs (female guts) or 30 mins (discs). They were then washed in PBS + 0.1% Triton X-100 (Euromedex 2000-C) (PBT). To remove the content of the gut lumen, the fixed guts were equilibrated 30 mins in PBS-50% Glycerol (Sigma-Aldrich G9012) and then washed in PBT to osmotically release gut contents. The primary antibody staining was performed overnight at 4°C in PBT. After washing with PBT 20 mins, 3 times, the secondary antibodies were incubated 3-4 hrs at room temperature (RT). After washing with PBT 2 times, nuclei were counterstained with DAPI in PBT for 20 mins (1µg/ml).

#### Discs

Adapted from a protocol provided by V. Loubiere (^75^ G. Cavalli lab). Wandering larvae were collected, and wing imaginal discs were quickly dissected at RT in 1× PBS before being fixed for 20 min in 4% paraformaldehyde. Imaginal discs were then permeabilized for 1 hour in 1× PBS + 0.5% Triton X-100 and blocked for 1 hour in 3% bovine serum albumin (BSA) PBTr (1× PBS + 0.025% Triton X-100). Then, primary antibodies were added in 1% BSA PBTr and incubated overnight at 4°C. The following day, discs were washed in PBTr, secondary antibodies were added and incubated for 2 hours at RT. After washing with PBTr, nuclei were counterstained with DAPI in PBTr for 20 mins (1µg/ml).

#### Antibodies

The antibodies used were: chicken anti-GFP (1:2000, Invitrogen #A10262), rabbit anti-DsRed (1:1000, Clontech #632496), mouse anti-RFP (Invitrogen (RF5R), rabbit anti-γH2Av (1:2000, Rockland 600-401-91), rabbit anti-Inx7^76^ (1:100, R. Bauer), guinea pig anti-Inx2^77^ (1:1000, G. Tanentzapf), rabbit anti-Inx2^78^ (1:1000, G. Ammer), rabbit anti-Inx3^79^ (1:70, R. Bauer), rabbit anti-Inx1^78^ (1:1000, G. Ammer), rabbit anti-Inx4^78^ (1:1000, G. Ammer), rabbit anti-Inx5^78^ (1:500, G. Ammer), rabbit anti-Inx6^78^ (1:1000, G. Ammer), rabbit anti-Inx8 serum^78^ (1:1000, G. Ammer), goat anti-βGal (1:200, F. Schweisguth), chicken anti-βGal (1:2000, Abcam ab9361), rabbit anti-PH3 (ser10, 1:2000, Merck Millipore #06-570), rabbit anti-RnrL^30^ (1:200, A. Arnaoutov), rabbit anti-DNK^80^ (1:20000, A. Lalouette).

The antibodies mouse anti-γH2Av (1:200, DSHB UNC93-5.2.1), mouse anti-Delta (1:1000, DSHB C594.9B), mouse anti-Arm (1:200, DSHB N2 7A1), mouse anti-Pros (1:2000, DSHB MR1A-c), and mouse anti-Cora (1:100, DSHB C615.16) were developed respectively by R. S. Hawley (Stowers Institute), S. Artavanis-Tsakonas (Harvard Medical School), E. Wieschaus (Princeton University), C.Q Doe (University of Oregon), and R. Fehon (University of Chicago), and were obtained from the Developmental Studies Hybridoma Bank, created by the NICHD of the NIH and maintained at The University of Iowa, Department of Biology, Iowa City, IA 52242.

The secondary antibodies were from Jackson Laboratory, raised in donkey with different fluorophores (DyLight 488, 549, 649), see table S3 for reference.

### 5-ethynyl-2 deoxyuridine (EdU) treatment and labelling

For the EdU incorporation experiment in wing discs, wandering larvae (late L3 stage) were collected and dissected in Schneider’s culture medium supplemented with L-glutamine (Sigma-Aldrich S0146) for 10mins. Then, EdU (Invitrogen by Thermo Fisher Scientific -C10340) was added to the dissection medium at a final concentration of 1µM, and the dissected wing discs were incubated for 30 mins at 25°C. After washing once quickly with PBS, the wing discs were fixed with PFA and stained with different antibodies following the procedure detailed above. The EdU Click-iT^TM^ reaction was then performed for 30 mins following the manufacturer’s protocol to label incorporated EdU with Alexa Fluor^TM^ 647 dye (Invitrogen by Thermo Fisher Scientific - C10340). After washing with PBTr, nuclei were counterstained with DAPI in PBTr for 20 mins (1µg/ml).

For the EdU incorporation experiment in midguts, adult *Drosophila* female flies were dissected in Schneider’s culture medium supplemented with L-glutamine (Sigma-Aldrich S0146) for 10 mins. Then, EdU (Invitrogen by Thermo Fisher Scientific - C10340) was added to the dissection medium at a final concentration of 10 µM and the dissected midguts were incubated for 3 hours at 25°C in a humid chamber. After washing once quickly with PBS, the midguts were fixed with PFA and stained with different antibodies following the procedure detailed above. The EdU Click-iT^TM^ reaction was then performed for 30 mins following the manufacturer’s protocol to label incorporated EdU with Alexa Fluor^TM^ 647 dye (Invitrogen by Thermo Fisher Scientific - C10639). After washing with PBT, nuclei were counterstained with DAPI in PBT for 20 mins (1µg/ml).

### *Ecc15* bacterial infection

First, for each conditions 15 adult female flies were mixed with 3 males in a tube and transferred at 29°C for 2 days to induce stem cell specific dnk KD and/or GFP expression. *Ecc15* bacteria treatment were conducted as previously published^81^, briefly, after 4 hours of starvation at 29°C flies were treated in tubes on filter paper soaked with a 1:1 mix of OD200 Ecc15 bacterial culture and 5% sucrose (Ecc15 infection) or a 1:1 mix of LB and 5% sucrose (no infection control). Proliferation response was assayed 15 hours after the treatment started.

### Microscopy and Image Analysis

Microscopy images were taken at the Cell and Tissue Imaging Platform – PICT-IBiSA of the UMR3215 of Institut Curie using the Zeiss confocal microscope LSM900 with a water 25x objective (whole discs), oil 40x objective (most of the gut images), Zeiss confocal microscope LSM780 with oil 25x objective (whole-disc images); Zeiss Apotome microscope 10X (whole-gut images). The images were processed, and the quantification performed using Fiji, unless another software is mentioned below.

#### PH3+ cells, stem cell loss and MARCM clone size

For visualizations and quantifications of stem cell loss, MARCM clone size (cell number per clones) and PH3+ cells in adult midguts, we used a 20x dry objective on the Leica epifluorescence microscope DM6000B to score each phenotype.

#### FUCCI and MARCM analysis in the gut

FUCCI and clonal (MARCM) experiments in the gut used Fiji/ImageJ^82^ for microscope image processing and quantification. Cell nuclei were manually outlined in Fiji for γH2Av intensity measures. For the FUCCI experiment, GFP and RFP intensities were also measured in the outlined nuclei, the ratio of GFP and RFP intensity and a visual confirmation allowed the attribution of cell cycle phases to individual stem cells.

For the clonal (MARCM) experiments, the cells were attributed a specific identity based on Prospero, Delta and DAPI staining. ISCs were identified with Delta staining, EEs with nuclear Prospero staining, ECs with a large nucleus (>20µm^2^), and EBs had small nuclei (<20µm^2^) without Delta or Pros. The limit of 20µm^2^ area for an EC nucleus was determined based on GFP staining in MyoAGal4 flies, in which GFP expression is driven specifically in ECs.

#### γH2Av mean intensity and RpA70-GFP max intensity measures in ISCs

For the measurement of γH2Av mean intensity and RpA70-GFP max intensity in ISCs, in the case of Delta^ts^ driven GFP or RFP expression: The detection and outline of ISC nuclei was performed automatically on Fiji from maximum projections of the confocal Z-stacks images with a macro that we implemented: the DAPI and GFP intensity were used, respectively, to automatically outline the cell nuclei and the ISC cell area. The intersection of both “masks” indicated stem cell nuclei. The intensity thresholds used for both channels were determined for each experiment as the value depends on varying staining conditions and microscope settings. Several rounds of smoothing on Fiji allowed the refinement of ISC nuclei outlines. Importantly, for the analysis, nuclei with area <5µm^2^ or >20µm^2^ (ECs or nuclei fusion due to the maximum projection) were not considered as they likely do not represent stem cells. From each outlined nuclei, we collected the mean γH2Av intensity and the maximum RpA70-GFP intensity, corresponding to the brightest foci in each nucleus. The data were then normalized to the mean value obtained in the Control conditions for each experimental replicate. The pictures for which the identification of stem cell nuclei was difficult (too many nuclei fused in the maximum projection, or bad signal for DAPI or GFP) were discarded from the analysis.

#### RedFlipout clones γH2Av distance analysis

We used CellProfiler (^83^) for the analysis of γH2Av intensity in the FlipOut clones in the wing discs. The CellProfiler analysis allows the semi-automatic detection with manual validation and correction of disc area based on DAPI intensity, and clone area based on RFP expression. From this, we measured the mean intensity in clonal (RFP+ DAPI+) and wild-type non-clonal (RFP-DAPI+) area for each wing disc analyzed. For each wing disc, CellProfiler also determined the approximate cell localization and nuclear area for single cells in the clones based on DAPI staining. For each identified cell, we extracted the position, γH2Av mean intensity in the area outlined and the shortest distance to the border of the clone (i.e. wild-type non-clonal tissue). The γH2Av intensity was then normalized to the mean γH2Av intensity found in the wild-type non-clonal area of the same wing disc, used as a baseline. Altogether it enabled the analysis of the relationship between DNA damage (γH2Av intensity) and the distance to the clone border. For each disc, we also measured the maximum distance of the identified cell in the RFP+ clone to the non-clonal part of the disc, as a proxy for the size of the clone.

#### Engrailed γH2Av measurements

In the experiment with *RnrL* RNAi with the *engrailed-Gal4* (*en-Gal4*) driver, both GFP and γH2Av intensity were measured along the antero-posterior line of the wing disc. The GFP intensity was used to determine the border of the engrailed domain for each measurement. For all the other experiments with *en-Gal4* driver, the mean γH2Av intensity was measured in an area of the pouch region of the engrailed domain (containing most of it) and compared with a similar area on the opposite side of the same wing disc, giving a “γH2Av intensity ratio” for each wing disc.

#### TwinSpots

For the TwinSpot experiment, the wing discs were analyzed and imaged at the LSM780 Zeiss confocal microscope, images of all the discs with DNA damage were taken, and representative images of all phenotypes were also taken.

#### RnrL, Inx2, and Inx7 quantification in the gut

For the RnrL, Inx2 or Inx7 quantification in the midgut, the cell types were identified based on Su(H)GBE-LacZ expression, Prospero staining, and DAPI staining (nuclear size). EB have a small nucleus and are βGal+ with a strong Armadillo (Arm) membrane staining, EEs were Pros+, young EC had a big nucleus (polyploid) and were also βGal+, the older ECs had a big nucleus and low Arm staining. ISC were identified with strong Arm membrane staining, basal localization, and absence of βGal. If a few puncta of Inx2 or Inx7 were found at the membrane of the identified cells, these were considered with partial Inx2 or Inx7 localization at the membrane.

#### Adult wing mounting was performed as described in^84^

### Data Processing and Statistics

Data Analysis was performed on Excel, Prism (version 9.0.1) or R (version 4.2.2) using the Rstudio interface (version 2022.07.02) and ggplot2 package (version 3.4.0). For all the figures with a boxplot, the center line is the median, box limits are upper and lower quartiles and whiskers extend to the observation points less than or more than 1.5x interquartile range. Initial training for the use of the ggplot2 package for data visualization was obtained by the U900 Bioinformatics unit of the Curie Institute. As indicated in the figures, the statistical significance was represented as follows: *: p < 0.05; **: p <0.01; ***: p<0.001; ****: p<0.0001.

---

**Figure S1.**
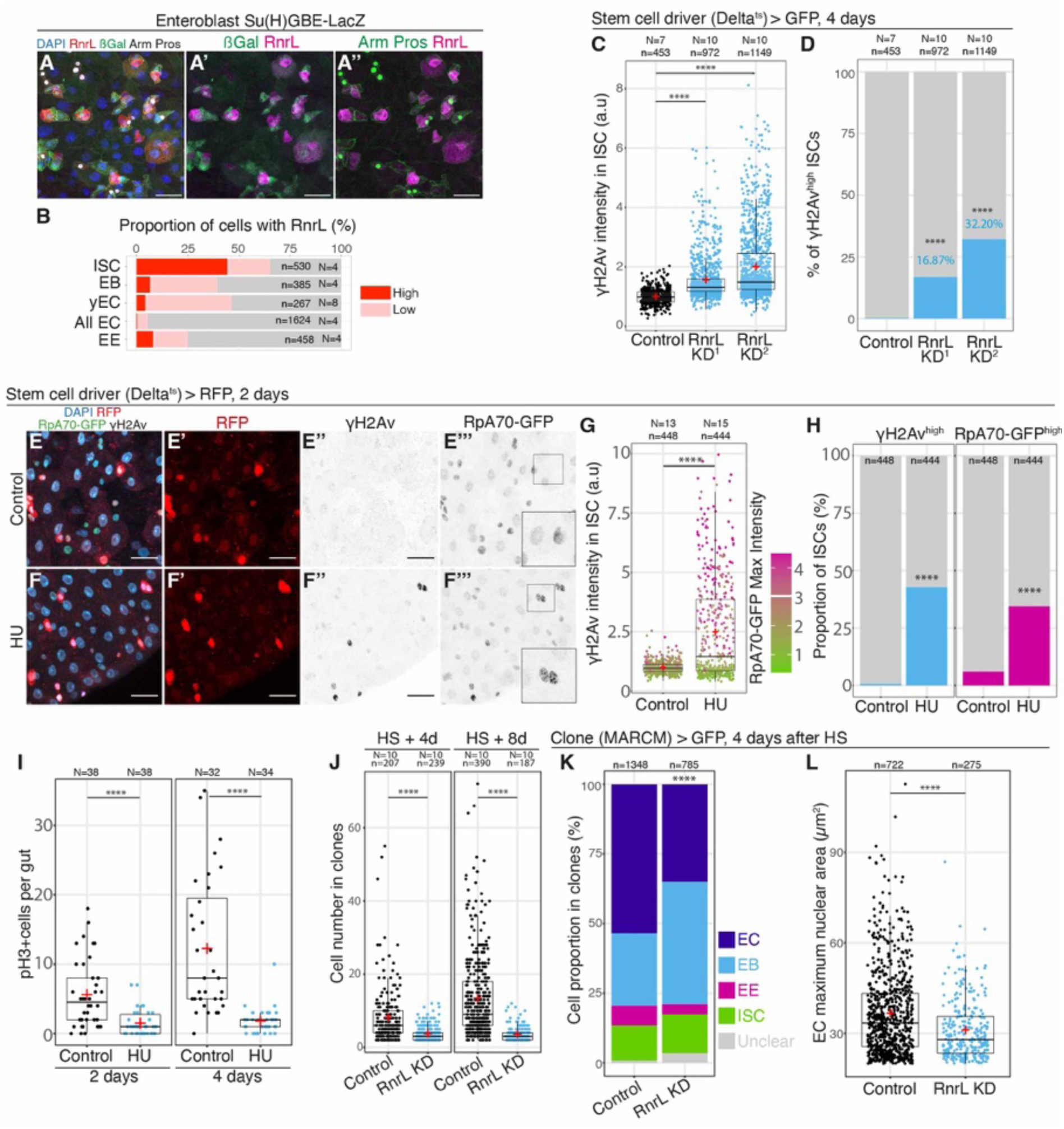
*RnrL* inhibition causes replication stress in midgut stem cells. **A-A ’’**. *Su(H)GBE-LacZ* midguts marked for RnrL, ßGal, Armadillo (membrane), and Prospero (nuclear). ßGal is expressed in EBs (diploid, small nuclei) and persists in newly produced young ECs (yECs, polyploid, big nuclei). ISCs are Arm+ ßGal-, EEs are Pros+, EBs are Arm+ ßGal+, ECs are characterized by large nuclei. Scale bar: 25µm. **B.** Percentage of midgut cells with high or low levels of RnrL protein expression as detected by immunostaining in A-A’’. yEC are newly produced ECs in which ßGal persists. **C.** γH2Av mean intensity in stem cell nuclei (Dl^ts^>GFP+) without or with *RnrL* RNAi for 4 days, two different RNAi lines were tested, normalized to the mean value in the Control. Welch’s t-test, two-sided, in comparison with the control. **D.** Proportion of ISCs (GFP+) with high levels of γH2Av (>2 of mean γH2Av intensity in C). Fisher’s test, one-sided. **E-F’’’.** *Delta^ts^*>RFP, RpA70-GFP female midguts fed with Control food (E-E’’’), or HU-containing food (F-F’’’) for 2 days; DAPI, RFP, γH2Av, and GFP are shown. Scale bar: 20µm. **G.** γH2Av mean intensity and RpA70-GFP max intensity in each ISC (RFP+) nucleus in Control or HU treated conditions, normalized to the mean value in the Control. Welch’s t-test, two-sided, in comparison with the control. **H.** Proportion of ISCs (RFP+) with high levels of γH2Av (>2 of mean γH2Av intensity in G) or RpA70-GFP (>3 of max RpA70-GFP intensity in G). Fisher’s test, two-sided. **I.** PH3+ cells per midgut in female flies fed with Control or HU containing food for 2 or 4 days. Welch’s t-test, two-sided. **J.** Cell number per clone (clone size) 4 days or 8 days after heat-shock in Control or *RnrL* RNAi conditions. Welch’s t-test, two-sided. **K.** Cell composition of MARCM>GFP clones with or without *RnrL* RNAi. Chi-squared test. The cells for which the Dl staining was not clearly positive or negative were classified as “Unclear”. **L.** EC nuclear size in MARCM>GFP clones with or without *RnrL* RNAi. Welch’s t-test, two-sided. Note that the size of EC nuclei was reduced, consistent with nucleotide depletion affecting EC polyploidization which requires replication. *: p < 0.05; **: p <0.01; ***: p<0.001; ****: p<0.0001. N = guts, n = cells. Red crosses = mean.

**Figure S2.**
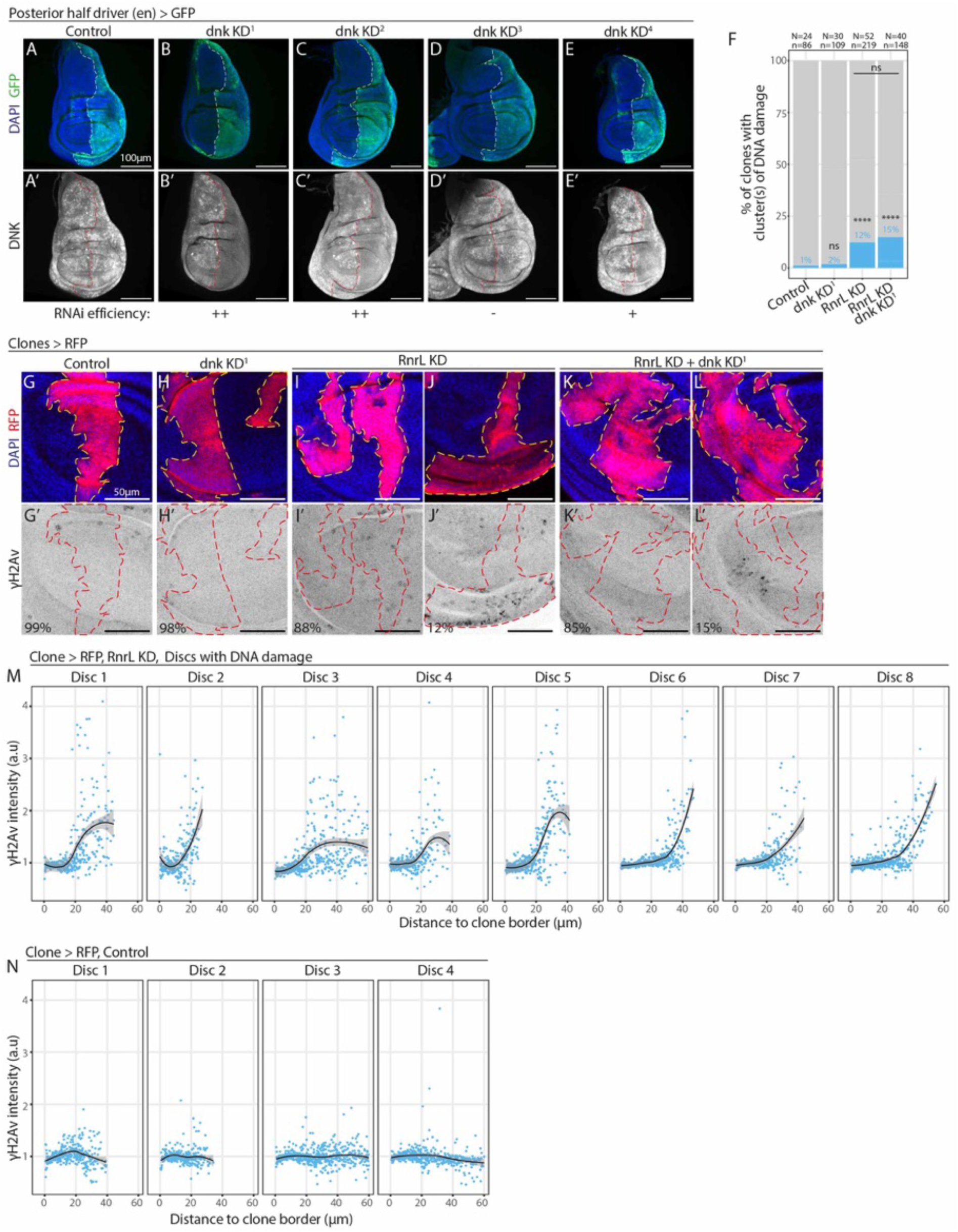
Investigating the salvage pathway DNA damage localization. **A-E’.** Test of *dnk* RNAi efficiency. *en>GFP* Control discs (w^1118^, A-A’), and discs expressing four different *dnk* RNAis(B-E’) marked for DAPI, GFP and DNK. The *dnk* KD 1 (B-B’) and 2 (C-C’) showed loss of DNK staining indicating an efficient knockdown of *dnk*. The *dnk* RNAi line 3 (D-D’) was not efficient while the 4^th^ RNAi (E-E’) had partial efficiency as DNK staining was reduced. The *dnk* KD^1^ line was used for the rest of the paper. Scale bar: 100µm. **F.** Percentage of clones displaying clusters of cells with DNA damage (γH2Av). N = wing discs, n= clones. Fisher’s test, one-sided. **G-L’.** hs- FlipOut>RFP clones in the late third instar larval wing disc, 4 days after induction in first instar larvae and stained for DAPI, RFP and γH2Av, with Control clones (G-G’), *dnk* RNAi (H-H’), *RnrL* RNAi (I-J’), and *dnk* & *RnrL* RNAi (K-L’). While most clones with *RnrL* RNAi or *dnk* & *RnrL* RNAi had no DNA damage (I-I’, K-K’), a minority of clones displayed clusters of cells with higher level of γH2Av (J-J’, L-L’). Scale bar: 50µm. **M-N.** Mean γH2Av intensity per cells in the clones relative to the distance to the wild-type surrounding tissue (clone border), in 8 different discs expressing *RnrL* RNAi in clones that displayed DNA damage (M), and 4 Control wild-type wing discs without DNA damage (N). For each wing disc, the black line is the LOESS curve of local regression (y ∼ x) and the gray area represents a 0.95 confidence interval. The value is normalized to the mean γH2Av intensity in wild-type non-clonal tissue (outside of the clones) for each wing discs. DNA damage was found 20-35µm from the clone border. **O-P’.** *en>GFP* Control discs (*w^11^*^18^, O-O’), and discs expressing *RnrL* RNAi (P-P’) marked for DAPI, GFP and RnrL. RnrL is lost throughout the engrailed (GFP+) domain in the *RnrL* KD condition. *: p < 0.05; **: p <0.01; ***: p<0.001; ****: p<0.0001.

**Figure S3.**
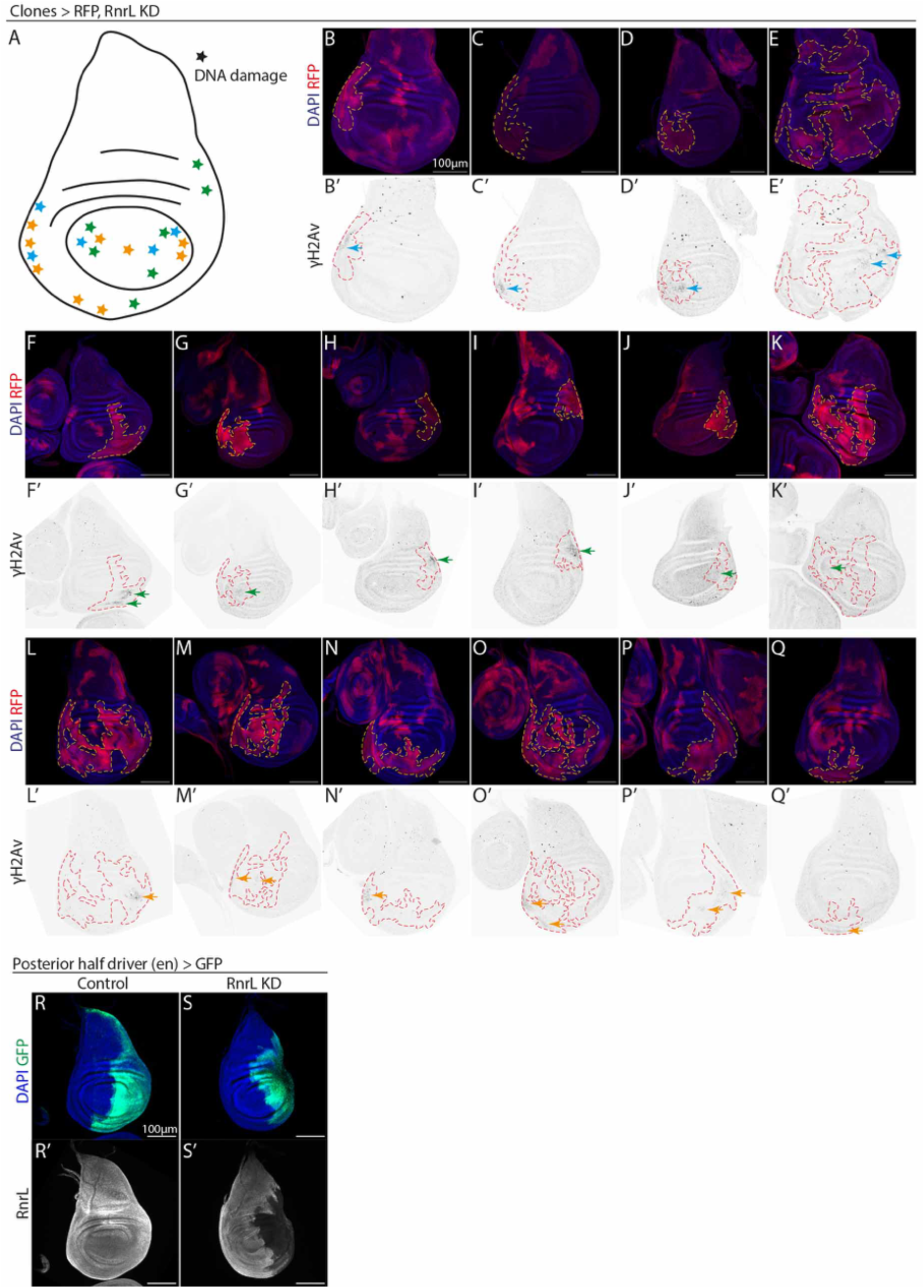
Position of clones with DNA damage in the wing disc. **A.** Schematic showing position of DNA damage within those clones having damage. Colors represent disctinct biological replicates. **B-Q’.** Discs with DNA damage (γH2Av), represented in A. **R-S’** RnrL protein expression in Control or *RnrL* KD conditions using en-Gal4.

**Figure S4.**
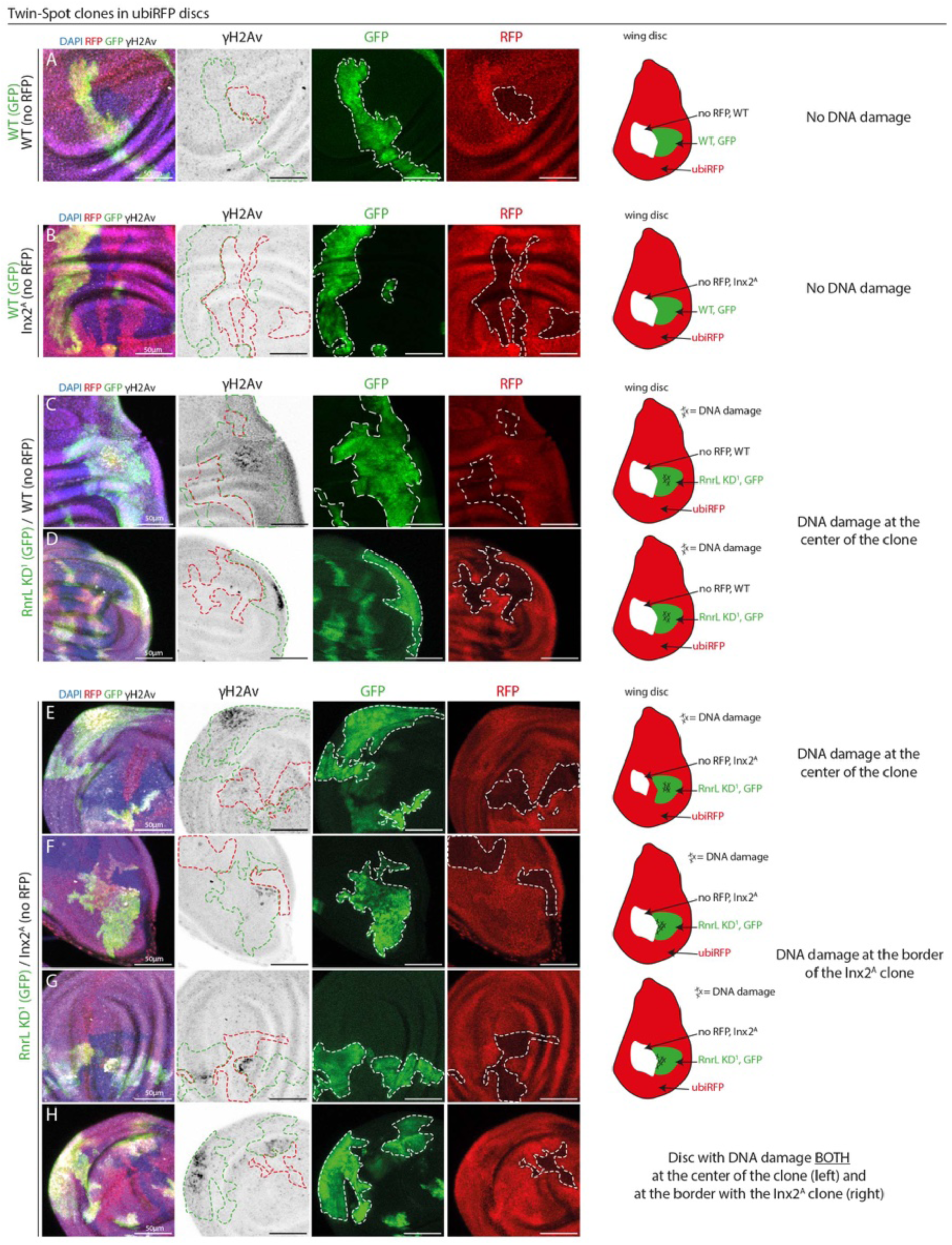
Twin spot-clones in the wing disc. **A-H.** Examples of twin-spot clones in the and stained for DAPI, GFP, RFP and γH2Av. As indicated in the schematic, while the tissue expresses RFP ubiquitously, upon clone induction, one clone loses the RFP expression and remains wild-type (WT; A, C-D) or becomes *Inx2^A^* homozygous mutant (B, E-H), and the twin-clone obtains Gal4/UAS driven expression of GFP with *RnrL* RNAi (C-H) or without *RnrL* RNAi (A, B), resulting in the corresponding genotype for each induced clones: WT/WT (A), WT/*Inx2^A^* (B), *RnrL* RNAi/WT (C, D) and *RnrL* RNAi/*Inx2^A^ (E-H)*. The controls A-B had no DNA damage.

**Figure S5.**
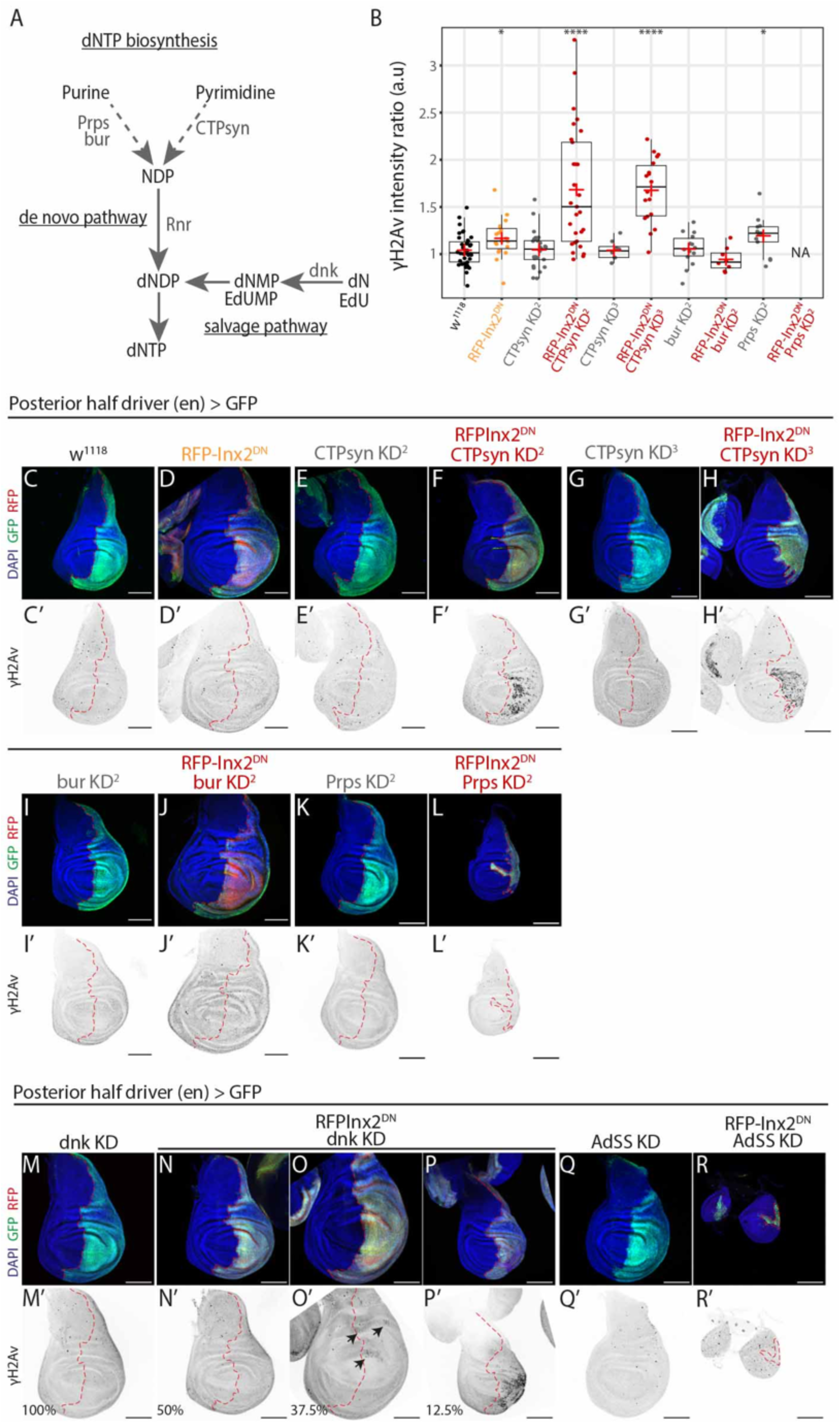
Nucleotide sharing through gap junctions in the wing disc epithelium. **A.** Simplified representation of the dNTP biosynthesis pathways and enzymes. EdU nucleoside analog (used in Figure 7) is processed by the salvage pathway. **B.** γH2Av mean intensity in the pouch region of the engrailed domain (posterior, GFP+) relative to the mean intensity in the pouch region of the non-engrailed domain (anterior, GFP-). Welch’s t-test, two-sided, in comparison with the control. **C-L’.** Late third instar larval *en>GFP* control discs (*wildtype*, C-C’), disc expressing a dominant negative form of Inx2 (RFPInx2^DN^, D-D’), *CTP synthetase* RNAi (E-E’, G-G’), *bur* RNAi (I-I’), *Prps* RNAi (K-K’), or a dominant negative form of Inx2 in combination with *CTP synthetase* RNAi (F-F’, H-H’), *bur* RNAi (J-J’) or *Prps* RNAi (L-L’), marked for γH2Av. Scale bar: 100µm. **M-P’.** *en>GFP* disc expressing *dnk* RNAi alone (M-M’) or in combination with a dominant negative form of Inx2 (RFPInx2^DN^, N-P’), marked for γH2Av, DAPI, GFP, RFP. While discs with *dnk* RNAi alone did not have DNA damage (N=15, M-M’), 50% of the discs with *dnk* RNAi and RFPInx2^DN^ displayed an increase in DNA damage, in small clusters of cells (37.5%, N=6/16, arrows in O-O’), or in most of the engrailed domain (12.5%, N=2/16, P-P’). **Q-R’.** AdSS (Adenylosuccinate Synthetase) is involved in ATP and dATP synthesis. *AdSS* KD alone appeared wild-type with respect to growth and DNA damage (Q-Q’). However, *AdSS* KD with concomitant inactivation of Inx2, showed a strong growth phenotype (R, R’). Scale bar: 100µm. *: p < 0.05; **: p <0.01; ***: p<0.001; ****: p<0.0001. N = discs. Red cross = mean

**Figure S6.**
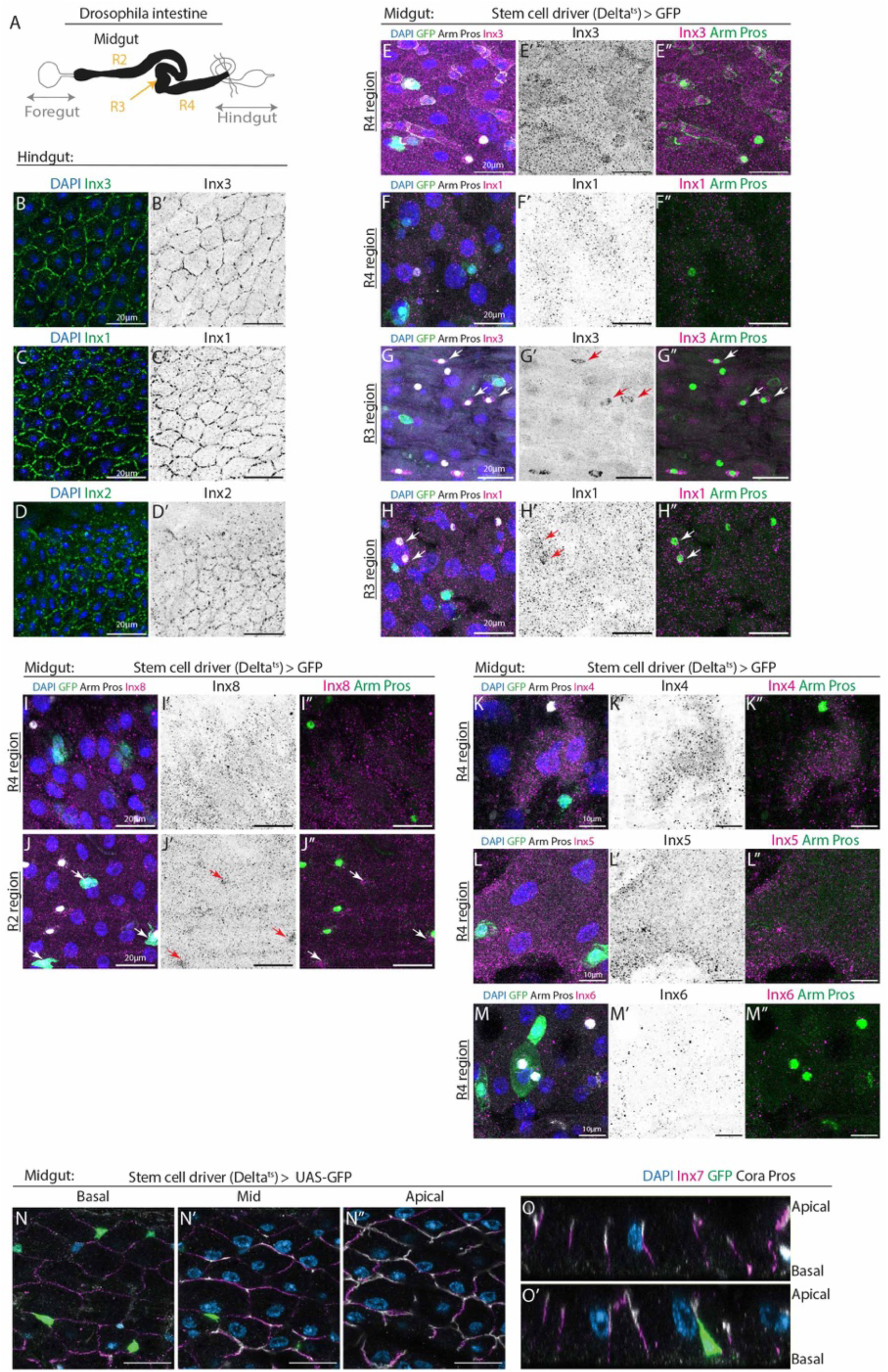
Innexin expression and localization in the *Drosophila* gut. **A.** Model of the D*rosophila* gut. **B-D’.** Hindgut tissue (adjacent to midgut): Membrane localization of Inx3 (B-B’), Inx1 (C-C’) and Inx2 (D-D’), consistent with formation of gap junction in this tissue. Scale bar: 20µm. **E-H’’’.** Midgut tissue: *Delta^ts^>GFP* female midguts stained for DAPI, GFP, Armadillo (membrane), Prospero (nuclear) and Inx3 (R4 region in E-E’’, R3 region in G-G’’) or Inx1 (R4 region in F-F’’, R3 region in H-H’’). *Delta^ts^*drives GFP expression in the ISCs, Prospero marks Enteroendocrine cells, and the ß-catenin Armadillo is enriched at the membrane of diploid cells (Intestinal stem cells and Enteroblasts). Inx3 and Inx1 appeared to form intracellular puncta in all the cells and was not enriched at any cell membranes throughout the midgut (E-F’’), with the exception of a small population of enteroendocrine (EE) cells in the middle midgut (R3 region) that had high levels of Inx3 and Inx1 throughout the cell (arrows, G-H’’). Scale bar: 20µm. **I-J’’.** *Delta^ts^>GFP* female midguts stained for DAPI, GFP, Armadillo (membrane), Prospero (nuclear) and Inx8. Inx8 appeared to form intracellular puncta in all the cells and was not enriched at any cell membranes (R4 region shown, I-I’’). Around 50% of intestinal stem cells (ISC) in the anterior midgut (R2 region) had higher levels of Inx8 in the cell (arrows, J-J’’) but not at the cell membrane. Scale bar: 20µm. **K-M’’.** *Delta^ts^>GFP* female midguts stained for DAPI, GFP, Armadillo (membrane), Prospero (nuclear) and Inx4, Inx5 or Inx6. Inx4, Inx5 and Inx6 were undetectable. Scale bar: 10µm. **N-O’.** *Delta^ts^>GFP* midguts marked for Inx7 for gap junctions and Coracle for septate junctions. Different views of the epithelium, basal (N), intermediate (N’) and apical (N’’), as well as orthogonal reconstitution from the Z-stack (O-O’). Inx2 and Inx7 proteins localized at the basal membranes below septate junctions in ECs. Scale bar: 20µm.

**Figure S7.**
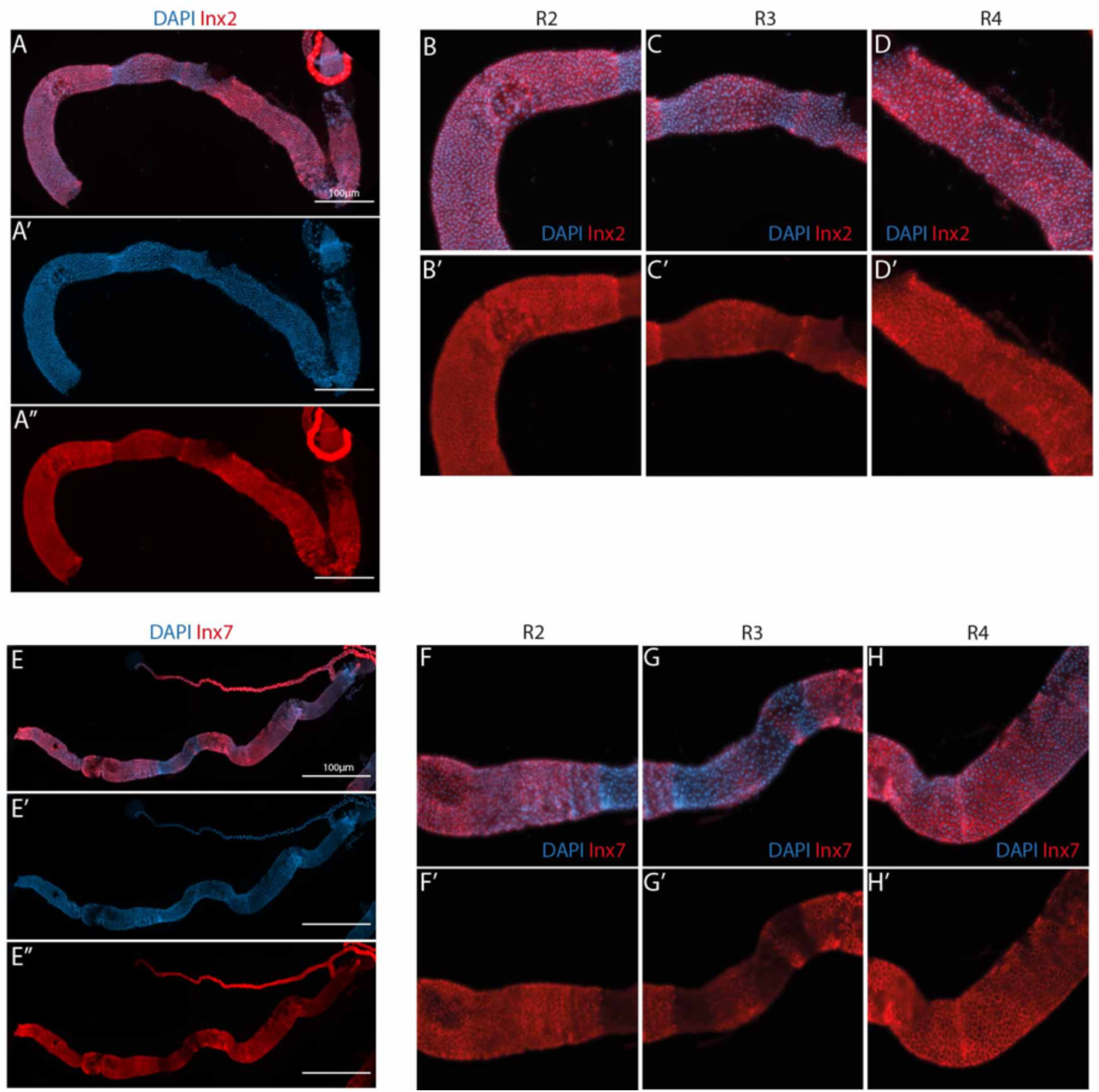
Regional differences in gap junction expression in the midgut. **A-D’.** Inx2 staining in whole midguts (A-A’’) and in close ups of anterior region (R2, B-B’), mid/copper cell region (R3, C-C’) and posterior (R4, D-D’). **E-H’.** Inx7 staining in whole midguts (E-E’’) and in close ups of anterior region (R2, F-F’), mid/copper cell region (R3, G-G’) and posterior region (R4, H-H’). Scale bars: 100µm. Inx2 and Inx7 were found to be highly expressed in R2 and R4 regions, expressed in R3, and absent in the R5 region of the midgut. Interestingly, different regions are separated by clear stripes lacking gap junction expression, likely separating different functional domains. A thin band of expression is found between R3 and R4 regions between two domains lacking expression.

**Figure S8.**
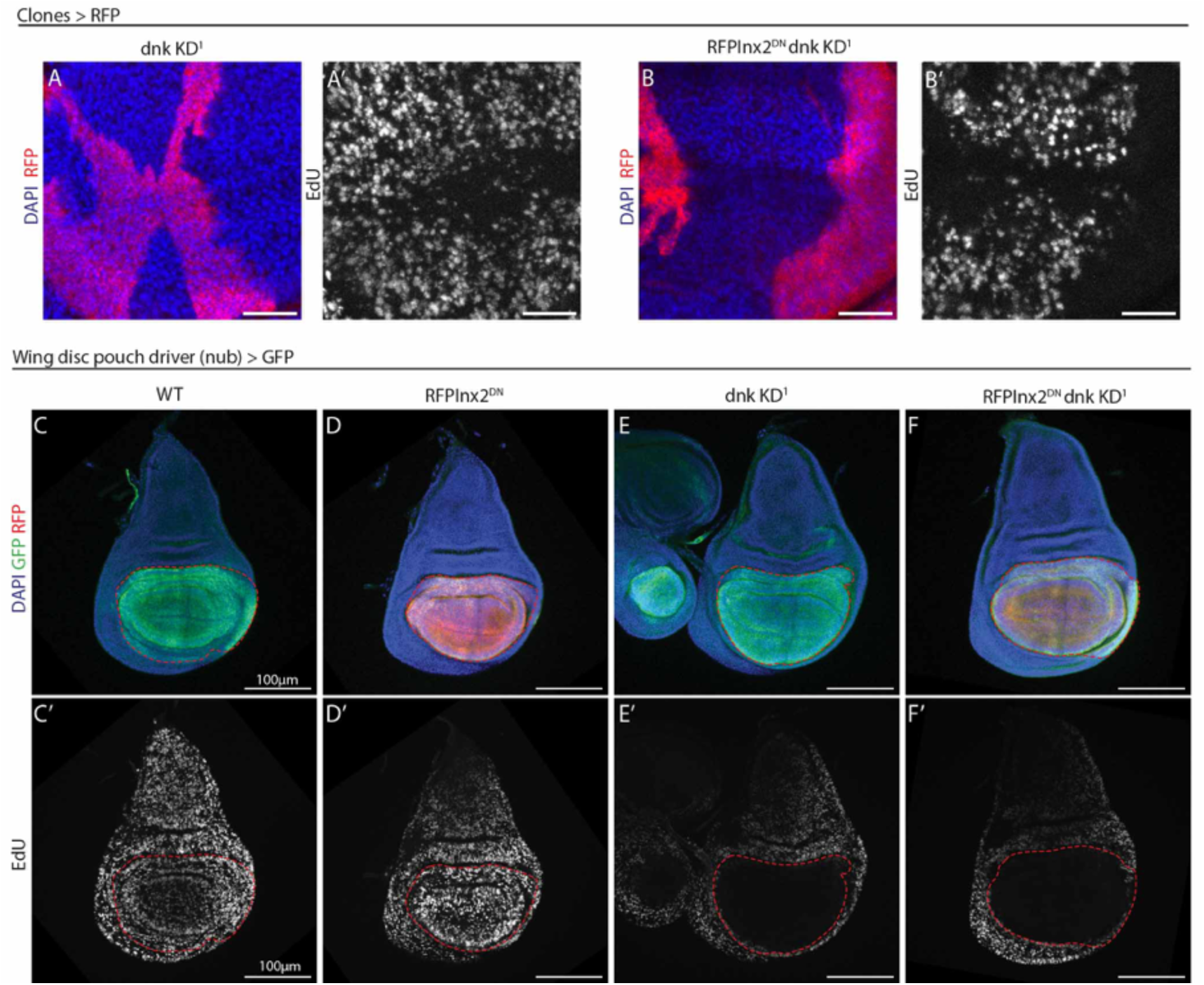
EdU nucleotide sharing assay in clones and the pouch region of the wing disc. **A-B ’.** hs-FlipOut>RFP clones in the late third instar larval wing disc, expressing *dnk* RNAi alone (A-A’) or in combination with a dominant negative form of Inx2 (RFPInx2^DN^, B-B’), 4 days after induction in first instar larvae and stained for DAPI, RFP and EdU labelling after 30 mins incorporation at the late L3 larval stage. Scale bar: 20µm **C-F’.** *nub>GFP* Control disc (*D-D’*), disc expressing a dominant negative form of Inx2 (RFPInx2^DN^, E-E’), dnk RNAi #1 (F-F’) or both (G-G’), labelled for EdU after 30 mins incorporation at the late L3 larval stage. When *dnk* is depleted in the whole pouch region, there is no gradient of EdU into the pouch, indicating the absence of transfer to the pouch from the neighboring tissue. Scale bar: 100µm.

**Table S1.** List of fly stocks used in the paper.

| Name | Genotype | Source-Reference |
| --- | --- | --- |
| RnrL KD-1 | y[1] sc[*] v[1];; P{y[+t7.7] v[+t1.8]=TRiP.HMC02351}attP2 | BL51418 |
| RnrL KD-2 | y[1] sc[*] v[1]; P{y[+t7.7] v[+t1.8]=TRiP.HMS02737}attP40 ; | BL44022 |
| Inx2 KD | :: UAS-Inx2 RNAi TRIP JF02446 | P. Spéder (formerly BL28915) |
| RFP-Inx2DN | ; pTRW-inx2 UAS-RFP-inx2 (RI1) ; | A. Brand/P. Spéder |
| CTPsyn KD-1 | y[1] v[1]; P{y[+t7.7] v[+t1.8]=TRiP.HMJ02094}attP40 ; | BL53378 |
| CTPsyn KD-2 | y[1] v[1];; P{y[+t7.7] v[+t1.8]=TRiP.HM04062}attP2 | BL31752 |
| CTPsyn KD-3 | y[1] v[1];; P{y[+t7.7] v[+t1.8]=TRiP.JF02214}attP2 | BL31924 |
| Prps KD-1 | y[1] sc[*] v[1] sev[21]; P{y[+t7.7] v[+t1.8]=TRiP.HMC05080}attP40 ; | BL60086 |
| Prps KD-2 | y[1] sc[*] v[1] sev[21];; P{y[+t7.7] v[+t1.8]=TRiP.GL00463}attP2/TM3, Sb[1] | BL35619 |
| dnk KD-1 | w[1118];; P{GD15092}v39137 | v39137 |
| dnk KD-2 | ; P{KK100376}VIE-260B ; | v103385 |
| dnk KD-3 | y[1] sc[*] v[1] sev[21]; P{y[+t7.7] v[+t1.8]=TRiP.HMC06148}attP40 ; | BL65886 |
| dnk KD-4 | y[1] sc[*] v[1] sev[21];; P{y[+t7.7] v[+t1.8]=TRiP.GL00100}attP2 | BL35219 |
| bur KD-1 | y[1] v[1]; P{y[+t7.7] v[+t1.8]=TRiP.HMJ22744}attP40 bur RNAi ; | BL60432 |
| bur KD-2 | y[1] v[1];; P{y[+t7.7] v[+t1.8]=TRiP.JF01503}attP2 | BL31055 |
| AdSS KD | y[1] sc[*] v[1] sev[21];; P{y[+t7.7] v[+t1.8]=TRiP.HMS00956}attP2 | BL 33993 |
| w<sup>1118</sup> | w[1118] ;; | M. McVey |
| EnGal4 GFP | w[*] ; P[En-Gal4], P[UAS-GFP]/CyO, w+, Tb ; | Y. Bellaiche, P. Leopold |
| Inx2A FRT19A | y[1] w[*] Inx2[A] P{ry[+t7.2]=neoFRT}19A/FM7c, P{w[+mC]=GAL4-Kr.C}DC1, P{w[+mC]=UAS-GFP.S65T}DC5, sn[+] | BL54481 |
| ΔGal4 | :: P[Delta-Gal4]/TM6 Tb | S. Hou |
| UAS-GFP | w[*]; P{w[+mC]=UAS-2xEGFP}AH2 ; | BL6874 |
| UAS-nlsRFP | w[1118]; P{w[+mC]=UAS-RFP.W}2 ; | BL30556 |
| Dlts GFP | w[*] ; P{w[+mC]=UAS-2xEGFP}AH2/(CyO) ; P[Delta-Gal4], tub-gal80ts/TM6B, Tb, Hu | This study |
| Dlts RFP | w[*] ; P{w[+mC]=UAS-RFP.W}2/ (CyO) ; P[Delta-Gal4], tub-gal80ts/TM6B, Tb, Hu | This study |
| Dlts FUCCI | w[*] ; UAS-GFP-E2F1<sub>1-230</sub>#32 UAS-mRFP1-CycB<sub>1-266</sub>#13 ; P[Delta-Gal4], tub-gal80ts/TM6B, Tb, Hu | This study |
| FUCCI | ; UAS-GFP-E2F1 , UAS-RFP-CycA-nls ; MKRS/TM6 | B. Edgar |
| RpA70-GFP | :: RPA-70::EGFP | A. M. Vales |
| TwinSpot driver | P{w[+mC]=Ubi-mRFP.nls}1, w[*], P{ry[+t7.2]=hsFLP}12 P{ry[+t7.2]=neoFRT}19A ; P{w[+mC]=Act5C-GAL4}25FO1, P{w[+mC]=UAS-GFP.U}2/CyO ; | This study (from BL42726, BL31406) |
| TwinSpot WT/WT | P{w[+mC]=tubP-GAL80}LL1 y[1] w[*] P{ry[+t7.2]=neoFRT}19A ;; | This study (from BL42726) |
| TwinSpot WT/Inx2A | P{w[+mC]=tubP-GAL80}LL1 y[1] w[*] Inx2[A] P{ry[+t7.2]=neoFRT}19A ;; | This study (from BL42726, BL54481) |
| TwinSpot RnrL KD/WT | P{w[+mC]=tubP-GAL80}LL1 y[1] w[*] P{ry[+t7.2]=neoFRT}19A ;; P{y[+t7.7] v[+t1.8]=TRiP.HMC02351}attP2 | This study (from BL42726, BL51418) |
| TwinSpot RnrL KD/Inx2A | P{w[+mC]=tubP-GAL80}LL1 y[1] w[*] Inx2[A] P{ry[+t7.2]=neoFRT}19A/FM7-GFP ;; P{y[+t7.7] v[+t1.8]=TRiP.HMC02351}attP2 | This study (from BL42726, BL54481, BL51418) |
| Su(H)GBE-LacZ | ;Su(H)GBE-LacZ | S. Bray |
| MARCM40<br>A | yw [hs-Flp]1.22 P[Tub-FRT-Gal4] P[UAS-nlsGFP]/(FM6) ; P{tubP-GAL80}LL10 P{neoFRT}40A/(CyO) ; | Bardin et al. 2010 (from BL5192) |
| FRT40A | ; {neoFRT}40A [mw]/ (CyO) ; | Bardin et al. 2010 |
| hs-FlipOut<br>RFP | hs-flp ;; Tub-FRT-Gal80-FRT-Gal4, UAS-mRFP/TM6b, Tb, Hu | Y. Bellaiche, R. Basto |
| MARCM19<br>A | p{ry[+] neoFR}19A, P{w[+] tubP-GAL80}L1, P{ry[+] hsFLP}1, w ; CyO/P{w[+] UAS-CD8:GFP}LL5 ; TM6, Tb, Hu / P{w[+] tubP-GAL4}LL7 | A. Gould |
| nub-Gal4 | w*; P{GawB}nubbin-AC-62; | P. Leopold |
| nubGal4<br>GFP | ; nubbin-GAL4, UAS:mCD8:GFP / CyO ; | Y. Bellaiche |
| Pdm2-Gal4 | ;; Pdm2-Gal4 TPK 841(Janelia R11F02) | P. Leopold |
| UAS-RnrL-RnrS | ;P{UAS-rnrL-P2A-rnrS}<sup>VK00018</sup>; | Arnaoutov et al., 2020<br>(A. Arnaoutov/M. Dasso) |

**Table S2.**
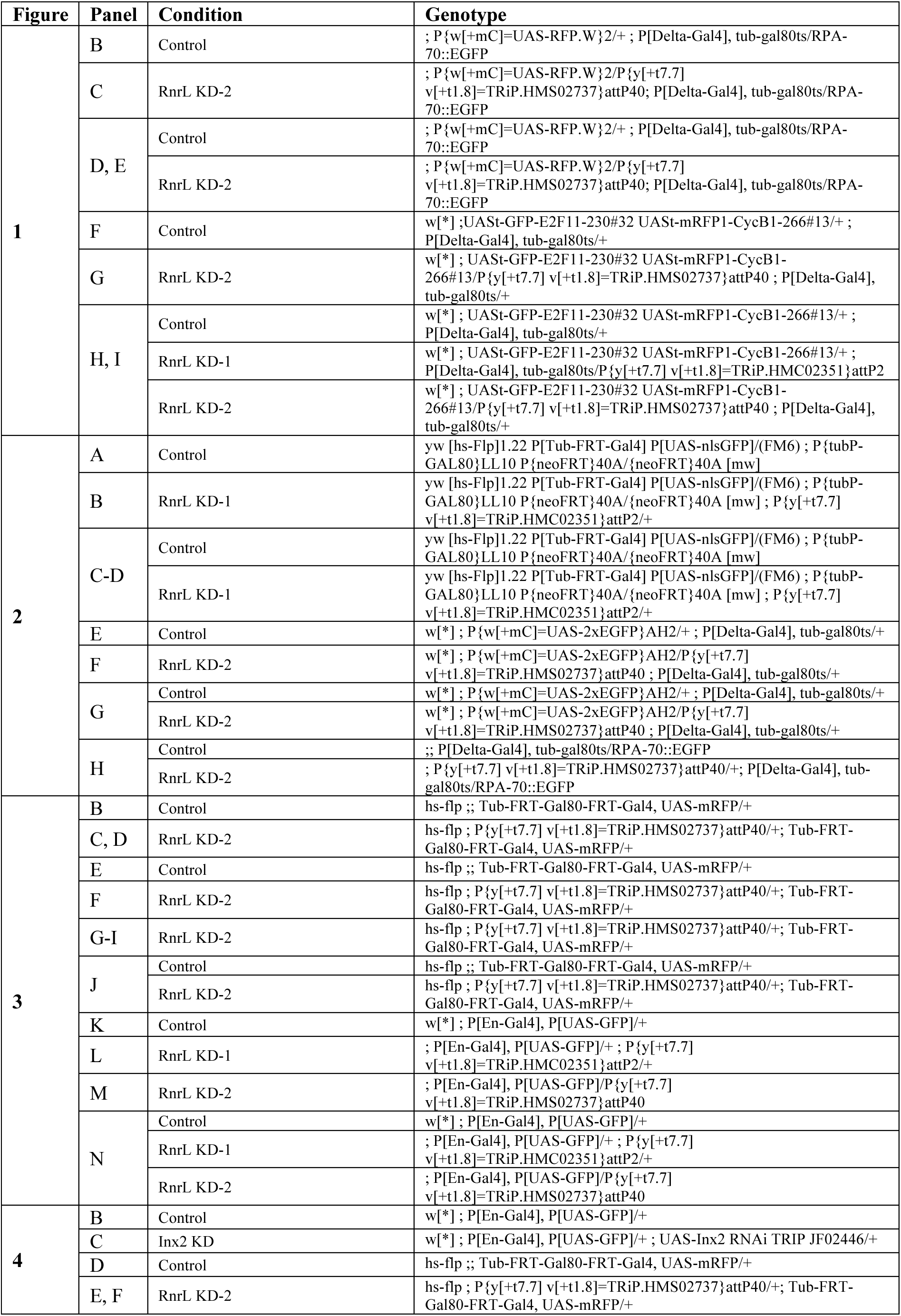
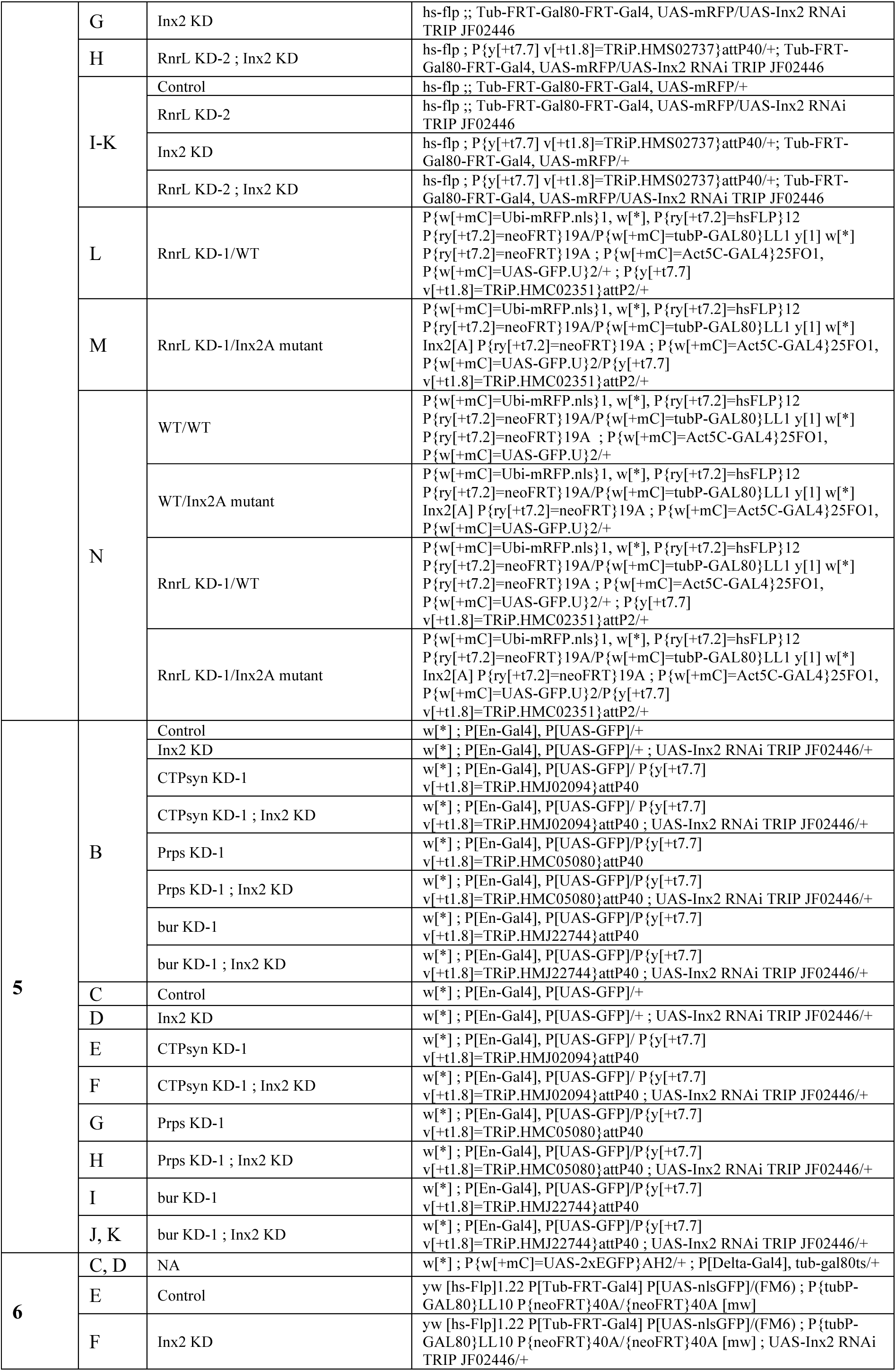
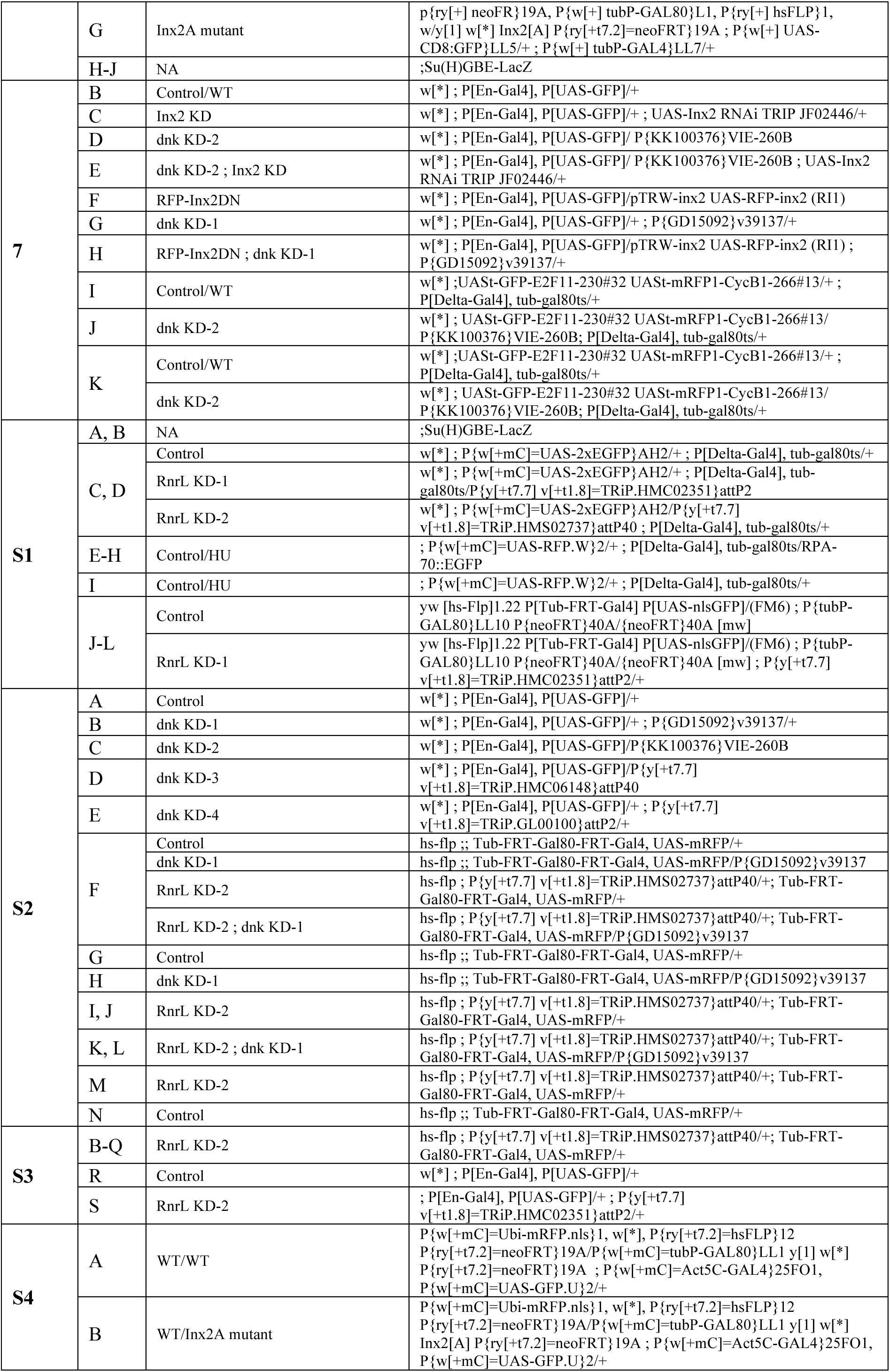
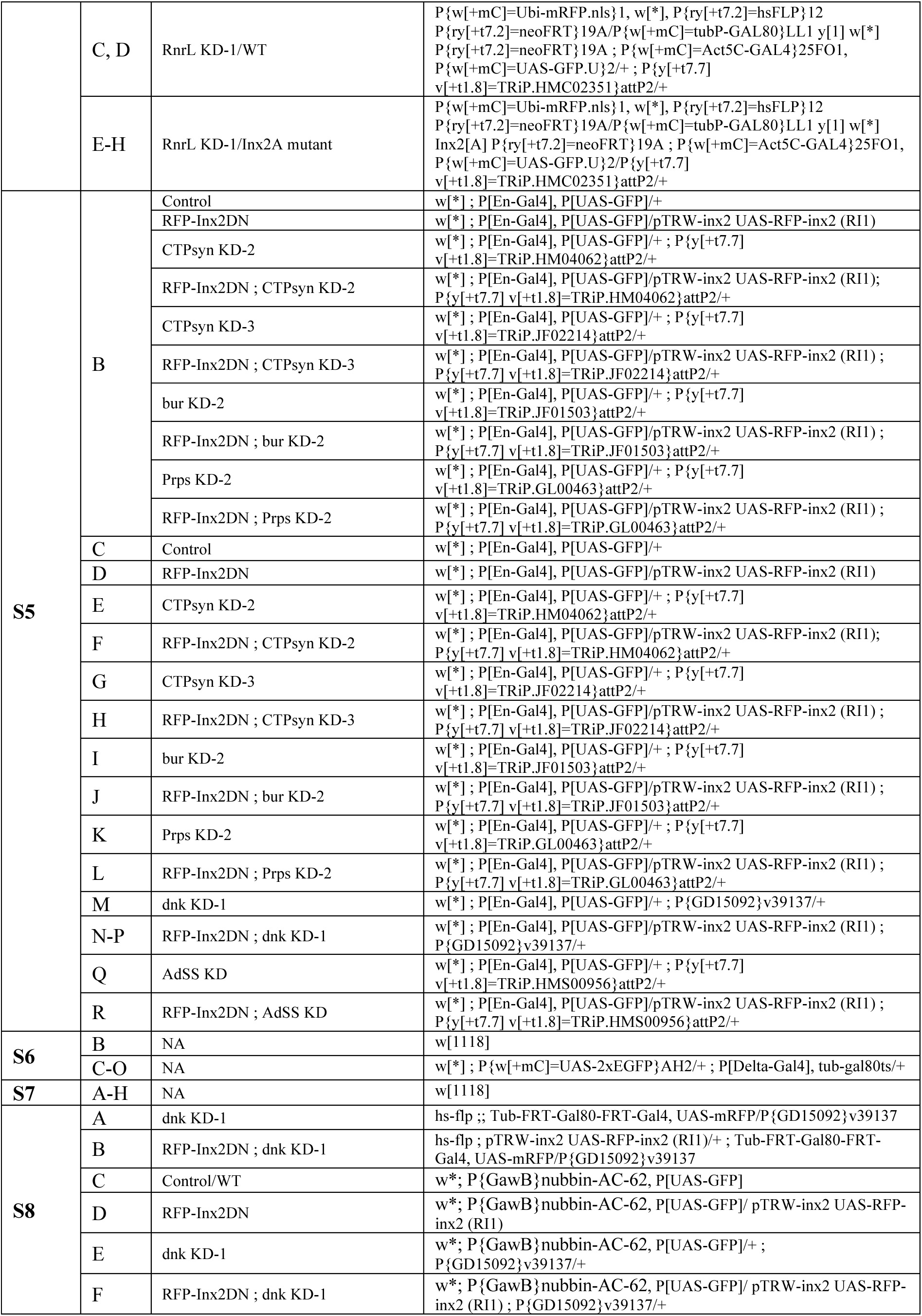
List of genotypes per figures.

**Table S3.** List of antibodies.

| **Antibody** | **Host** | **Source (Reference)** | **Dilution used** |
| --- | --- | --- | --- |
| GFP | Chicken | Invitrogen A10262 | 1:2000 |
| RFP | Mouse | Invitrogen (RF5R) | 1:2000 |
| RFP | Rabbit | Takara Living Colors 632496 | 1:1000 |
| γH2Av | Rabbit | Rockland 600-401-914 | 1:2000 |
| γH2Av | Mouse | DSHB UNC93-5.2.1 | 1:200 |
| RnrL | Rabbit | Arnaoutov et al., 2020 (A. Arnaoutov/M. Dasso) | 1:200 (gut), 1:100 (disc) |
| DNK | Rabbit | Knecht et al., 2000 (A. Lalouette) | 1:20000 |
| Armadillo | Mouse | DSHB N2 7A1 / F. Schweisguth | 1:200 |
| Delta | Mouse | DSHB C594.9ba | 1:1000 |
| Prospero | Mouse | DSHB MR1A-c | 1:2000 |
| Bgal | Goat | Biogenesis 466-1409 / F. Schweisguth | 1:200 |
| Bgal | Chicken | Abcam ab9361 | 1:2000 |
| Inx1 | Rabbit | Ammer et al., 2022 (G. Ammer) | 1:1000 |
| Inx2 | Rabbit | Ammer et al., 2022 (G. Ammer) | 1:1000 |
| Inx2 | Guinea Pig | Smendziuk et al., 2015 (G. Tanentzapf) | 1:500 |
| Inx3 | Rabbit | Bohrmann & Zimmermann, 2008 (R. Bauer) | 1:70 |
| Inx4 | Rabbit | Ammer et al., 2022 (G. Ammer) | 1:1000 |
| Inx5 | Rabbit | Ammer et al., 2022 (G. Ammer) | 1:500 |
| Inx6 | Rabbit | Ammer et al., 2022 (G. Ammer) | 1:1000 |
| Inx7 | Rabbit | Ostrowski et al., 2008 (R. Bauer) | 1:100 |
| Inx8 - serum | Rabbit | Ammer et al., 2022 (G. Ammer) | 1:1000 |
| Coracle | Mouse | DSHB C615.16 | 1:100 |
| pH3 | Rabbit | Merck Millipore 06-570 | 1:2000 |
| Cleaved Drosophila Dcp-1 (Asp216) | Rabbit | Cell Signaling 9578 | 1:100 |
| anti-Chicken DyLight-488 | Donkey | Jackson 703-486-155 | 1:2000 |
| anti-Goat DyLight-488 | Donkey | Jackson 705-486-147 | 1:2000 |
| anti-Guinea Pig DyLight-549 | Donkey | Jackson 706-506-148 | 1:2000 |
| anti-Guinea Pig DyLight-649 | Donkey | Jackson 706-496-148 | 1:2000 |
| anti-Mouse DyLight-488 | Donkey | Jackson 715-486-150 | 1:2000 |
| anti-Mouse DyLight-549 | Donkey | Jackson 715-506-150 | 1:2000 |
| anti-Mouse DyLight-649 | Donkey | Jackson 715-496-150 | 1:2000 |
| anti-Rabbit DyLight-488 | Donkey | Jackson 711-486-152 | 1:2000 |
| anti-Rabbit DyLight-549 | Donkey | Jackson 711-506-152 | 1:2000 |
| anti-Rabbit DyLight-649 | Donkey | Jackson 711-496-152 | 1:2000 |

## Notes

### Competing Interest Statement

The authors have declared no competing interest.

### Summary of Updates

We have added several, addition important experiments including a full characterization of location all of the fly Innexin proteins in the midgut and an assessment of EdU uptake into ISCs with expression dnk RNAi. In addition, we have extended our introduction and discussion.

## References

1. Techer, H., Koundrioukoff, S., Nicolas, A., and Debatisse, M. (2017). The impact of replication stress on replication dynamics and DNA damage in vertebrate cells. Nat Rev Genet 18, 535–550. 10.1038/nrg.2017.46.

2. Buj, R., and Aird, K.M. (2018). Deoxyribonucleotide Triphosphate Metabolism in Cancer and Metabolic Disease. Front Endocrinol (Lausanne) 9, 177. 10.3389/fendo.2018.00177.

3. 3. da Costa, A., Chowdhury, D., Shapiro, G.I., D’Andrea, A.D., and Konstantinopoulos, P.A. (2023). Targeting replication stress in cancer therapy. Nat Rev Drug Discov 22, 38–58. 10.1038/s41573-022-00558-5.

4. Mullen, N.J., and Singh, P.K. (2023). Nucleotide metabolism: a pan-cancer metabolic dependency. Nature Reviews Cancer. 10.1038/s41568-023-00557-7.

5. Zhu, J., and Thompson, C.B. (2019). Metabolic regulation of cell growth and proliferation. Nat Rev Mol Cell Biol 20, 436–450. 10.1038/s41580-019-0123-5.

6. Ali, E.S., and Ben-Sahra, I. (2023). Regulation of nucleotide metabolism in cancers and immune disorders. Trends Cell Biol. 10.1016/j.tcb.2023.03.003.

7. Toy, G., Austin, W.R., Liao, H.I., Cheng, D., Singh, A., Campbell, D.O., Ishikawa, T.O., Lehmann, L.W., Satyamurthy, N., Phelps, M.E., et al. (2010). Requirement for deoxycytidine kinase in T and B lymphocyte development. Proc Natl Acad Sci U S A 107, 5551–5556. 10.1073/pnas.0913900107.

8. Dobrovolsky, V.N., Bucci, T., Heflich, R.H., Desjardins, J., and Richardson, F.C. (2003). Mice deficient for cytosolic thymidine kinase gene develop fatal kidney disease. Mol Genet Metab 78, 1–10. 10.1016/s1096-7192(02)00224-x.

9. 9. Tran, P., Wanrooij, P.H., Lorenzon, P., Sharma, S., Thelander, L., Nilsson, A.K., Olofsson, A.K., Medini, P., von Hofsten, J., Stal, P., and Chabes, A. (2019). De novo dNTP production is essential for normal postnatal murine heart development. J Biol Chem 294, 15889–15897. 10.1074/jbc.RA119.009492.

10. Tran, D.H., Kim, D., Kesavan, R., Brown, H., Dey, T., Soflaee, M.H., Vu, H.S., Tasdogan, A., Guo, J., Bezwada, D., et al. (2024). De novo and salvage purine synthesis pathways across tissues and tumors. Cell 187, 3602–3618 e3620. 10.1016/j.cell.2024.05.011.

11. Wu, Z., Bezwada, D., Cai, F., Harris, R.C., Ko, B., Sondhi, V., Pan, C., Vu, H.S., Nguyen, P.T., Faubert, B., et al. (2024). Electron transport chain inhibition increases cellular dependence on purine transport and salvage. Cell Metab 36, 1504–1520 e1509. 10.1016/j.cmet.2024.05.014.

12. Stillman, B. (2013). Deoxynucleoside triphosphate (dNTP) synthesis and destruction regulate the replication of both cell and virus genomes. Proc Natl Acad Sci U S A 110, 14120–14121. 10.1073/pnas.1312901110.

13. Nordlund, P., and Reichard, P. (2006). Ribonucleotide reductases. Annu Rev Biochem 75, 681–706. 10.1146/annurev.biochem.75.103004.142443.

14. Robitaille, A.M., Christen, S., Shimobayashi, M., Cornu, M., Fava, L.L., Moes, S., Prescianotto-Baschong, C., Sauer, U., Jenoe, P., and Hall, M.N. (2013). Quantitative phosphoproteomics reveal mTORC1 activates de novo pyrimidine synthesis. Science 339, 1320–1323. 10.1126/science.1228771.

15. Ben-Sahra, I., Howell, J.J., Asara, J.M., and Manning, B.D. (2013). Stimulation of de novo pyrimidine synthesis by growth signaling through mTOR and S6K1. Science 339, 1323–1328. 10.1126/science.1228792.

16. French, J.B., Jones, S.A., Deng, H., Pedley, A.M., Kim, D., Chan, C.Y., Hu, H., Pugh, R.J., Zhao, H., Zhang, Y., et al. (2016). Spatial colocalization and functional link of purinosomes with mitochondria. Science 351, 733–737. 10.1126/science.aac6054.

17. Ben-Sahra, I., Hoxhaj, G., Ricoult, S.J.H., Asara, J.M., and Manning, B.D. (2016). mTORC1 induces purine synthesis through control of the mitochondrial tetrahydrofolate cycle. Science 351, 728–733. 10.1126/science.aad0489.

18. Subak-Sharpe, H., Burk, R.R., and Pitts, J.D. (1969). Metabolic co-operation between biochemically marked mammalian cells in tissue culture. J Cell Sci 4, 353–367. 10.1242/jcs.4.2.353.

19. Subak-Sharpe, I., Burk, R.R. and Pitts, J.D. (1966). Metabolic co-operation by cell to cell transfer between genetically different mammalian cells in tissue culture. Heredity 21, 342. 10.1038/hdy.1966.35.

20. Gilula, N.B., Reeves, O.R., and Steinbach, A. (1972). Metabolic coupling, ionic coupling and cell contacts. Nature 235, 262–265. 10.1038/235262a0.

21. Azarnia, R., Michalke, W., and Loewenstein, W.R. (1972). Intercellular communication and tissue growth. VI. Failure of exchange of endogeneous molecules between cancer cells with defective junctions and noncancerous cells. J Membr Biol 10, 247–258. 10.1007/BF01867858.

22. Cox, R.P., Krauss, M.R., Balis, M.E., and Dancis, J. (1974). Metabolic cooperation in cell culture: studies of the mechanisms of cell interaction. J Cell Physiol 84, 237–252. 10.1002/jcp.1040840210.

23. 23. Riddiford, N., Siudeja, K., van den Beek, M., Boumard, B., and Bardin, A.J. (2021). Evolution and genomic signatures of spontaneous somatic mutation in Drosophila intestinal stem cells. Genome Res 31, 1419–1432. 10.1101/gr.268441.120.

24. Siudeja, K., Nassari, S., Gervais, L., Skorski, P., Lameiras, S., Stolfa, D., Zande, M., Bernard, V., Rio Frio, T., and Bardin, A.J. (2015). Frequent Somatic Mutation in Adult Intestinal Stem Cells Drives Neoplasia and Genetic Mosaicism during Aging. Cell stem cell 17, 663–674. 10.1016/j.stem.2015.09.016.

25. Alvarez, S., Diaz, M., Flach, J., Rodriguez-Acebes, S., Lopez-Contreras, A.J., Martinez, D., Canamero, M., Fernandez-Capetillo, O., Isern, J., Passegue, E., and Mendez, J. (2015). Replication stress caused by low MCM expression limits fetal erythropoiesis and hematopoietic stem cell functionality. Nat Commun 6, 8548. 10.1038/ncomms9548.

26. Flach, J., Bakker, S.T., Mohrin, M., Conroy, P.C., Pietras, E.M., Reynaud, D., Alvarez, S., Diolaiti, M.E., Ugarte, F., Forsberg, E.C., et al. (2014). Replication stress is a potent driver of functional decline in ageing haematopoietic stem cells. Nature. doi:10.1038/nature13619.

27. Beerman, I., Bock, C., Garrison, B.S., Smith, Z.D., Gu, H., Meissner, A., and Rossi, D.J. (2013). Proliferation-dependent alterations of the DNA methylation landscape underlie hematopoietic stem cell aging. Cell stem cell 12, 413–425. 10.1016/j.stem.2013.01.017.

28. Boumard, B., and Bardin, A.J. (2021). An amuse-bouche of stem cell regulation: Underlying principles and mechanisms from adult Drosophila intestinal stem cells. Curr Opin Cell Biol 73, 58–68. 10.1016/j.ceb.2021.05.007.

29. Miguel-Aliaga, I., Jasper, H., and Lemaitre, B. (2018). Anatomy and Physiology of the Digestive Tract of Drosophila melanogaster. Genetics 210, 357–396. 10.1534/genetics.118.300224.

30. Arnaoutov, A., Lee, H., Plevock Haase, K., Aksenova, V., Jarnik, M., Oliver, B., Serpe, M., and Dasso, M. (2020). IRBIT Directs Differentiation of Intestinal Stem Cell Progeny to Maintain Tissue Homeostasis. iScience 23, 100954. 10.1016/j.isci.2020.100954.

31. 31. Zielke, N., Korzelius, J., van Straaten, M., Bender, K., Schuhknecht, G.F., Dutta, D., Xiang, J., and Edgar, B.A. (2014). Fly-FUCCI: A Versatile Tool for Studying Cell Proliferation in Complex Tissues. Cell Rep 7, 588–598. 10.1016/j.celrep.2014.03.020.

32. Syrjanen, J., Michalski, K., Kawate, T., and Furukawa, H. (2021). On the molecular nature of large-pore channels. J Mol Biol 433, 166994. 10.1016/j.jmb.2021.166994.

33. Bauer, R., Loer, B., Ostrowski, K., Martini, J., Weimbs, A., Lechner, H., and Hoch, M. (2005). Intercellular communication: the Drosophila innexin multiprotein family of gap junction proteins. Chem Biol 12, 515–526. 10.1016/j.chembiol.2005.02.013.

34. Sanchez, A., Castro, C., Flores, D.L., Gutierrez, E., and Baldi, P. (2019). Gap Junction Channels of Innexins and Connexins: Relations and Computational Perspectives. Int J Mol Sci 20. 10.3390/ijms20102476.

35. Ayukawa, T., Matsumoto, K., Ishikawa, H.O., Ishio, A., Yamakawa, T., Aoyama, N., Suzuki, T., and Matsuno, K. (2012). Rescue of Notch signaling in cells incapable of GDP-L-fucose synthesis by gap junction transfer of GDP-L-fucose in Drosophila. Proceedings of the National Academy of Sciences of the United States of America 109, 15318–15323. 10.1073/pnas.1202369109.

36. Dutta, D., Dobson, A.J., Houtz, P.L., Glasser, C., Revah, J., Korzelius, J., Patel, P.H., Edgar, B.A., and Buchon, N. (2015). Regional Cell-Specific Transcriptome Mapping Reveals Regulatory Complexity in the Adult Drosophila Midgut. Cell Rep 12, 346–358. 10.1016/j.celrep.2015.06.009.

37. Kim, A.A., Nguyen, A., Marchetti, M., Du, X., Montell, D.J., Pruitt, B.L., and O’Brien, L.E. (2022). Independently paced Ca2+ oscillations in progenitor and differentiated cells in an ex vivo epithelial organ. J Cell Sci 135. 10.1242/jcs.260249.

38. Petsakou, A., Liu, Y., Liu, Y., Comjean, A., Hu, Y., and Perrimon, N. (2023). Cholinergic neurons trigger epithelial Ca(2+) currents to heal the gut. Nature 623, 122–131. 10.1038/s41586-023-06627-y.

39. Lane, A.N., and Fan, T.W. (2015). Regulation of mammalian nucleotide metabolism and biosynthesis. Nucleic Acids Res 43, 2466–2485. 10.1093/nar/gkv047.

40. Diehl, F.F., Sapp, K.M., and Vander Heiden, M.G. (2023). The bidirectional relationship between metabolism and cell cycle control. Trends Cell Biol. 10.1016/j.tcb.2023.05.012.

41. Song, Y., Marmion, R.A., Park, J.O., Biswas, D., Rabinowitz, J.D., and Shvartsman, S.Y. (2017). Dynamic Control of dNTP Synthesis in Early Embryos. Dev Cell 42, 301–308 e303. 10.1016/j.devcel.2017.06.013.

42. Innis, M.A., Beers, T.R., and Craig, S.P. (1976). Mitochondrial regulation in sea urchins. I. Mitochondrial ultrastructure transformations and changes in the ADP:ATP ratio at fertilization. Exp Cell Res 98, 47–56. 10.1016/0014-4827(76)90461-4.

43. Dutta, S., Djabrayan, N.J., Smits, C.M., Rowley, C.W., and Shvartsman, S.Y. (2020). Excess dNTPs Trigger Oscillatory Surface Flow in the Early Drosophila Embryo. Biophys J 118, 2349–2353. 10.1016/j.bpj.2020.03.010.

44. Liu, B., Winkler, F., Herde, M., Witte, C.P., and Grosshans, J. (2019). A Link between Deoxyribonucleotide Metabolites and Embryonic Cell-Cycle Control. Curr Biol 29, 1187–1192 e1183. 10.1016/j.cub.2019.02.021.

45. Djabrayan, N.J., Smits, C.M., Krajnc, M., Stern, T., Yamada, S., Lemon, W.C., Keller, P.J., Rushlow, C.A., and Shvartsman, S.Y. (2019). Metabolic Regulation of Developmental Cell Cycles and Zygotic Transcription. Curr Biol 29, 1193–1198 e1195. 10.1016/j.cub.2019.02.028.

46. Loewenstein, W.R. (1979). Junctional intercellular communication and the control of growth. Biochim Biophys Acta 560, 1–65. 10.1016/0304-419x(79)90002-7.

47. Kanno, Y., and Loewenstein, W.R. (1964). Intercellular Diffusion. Science 143, 959–960. 10.1126/science.143.3609.959.

48. Kanno, Y., and Loewenstein, W.R. (1964). Low-Resistance Coupling between Gland Cells. Some Observations on Intercellular Contact Membranes and Intercellular Space. Nature 201, 194–195. 10.1038/201194a0.

49. Lo, C.W., and Gilula, N.B. (1979). Gap junctional communication in the post-implantation mouse embryo. Cell 18, 411–422. 10.1016/0092-8674(79)90060-6.

50. Kam, E., Melville, L., and Pitts, J.D. (1986). Patterns of junctional communication in skin. J Invest Dermatol 87, 748–753. 10.1111/1523-1747.ep12456937.

51. Kam, E., and Hodgins, M.B. (1992). Communication compartments in hair follicles and their implication in differentiative control. Development 114, 389–393. 10.1242/dev.114.2.389.

52. Pitts, J., Kam, E., Melville, L., and Watt, F.M. (1987). Patterns of junctional communication in animal tissues. Ciba Found Symp 125, 140–153. 10.1002/9780470513408.ch9.

53. Fraser, S.E., and Bryant, P.J. (1985). Patterns of dye coupling in the imaginal wing disk of Drosophila melanogaster. Nature 317, 533–536. 10.1038/317533a0.

54. Weir, M.P., and Lo, C.W. (1984). Gap-junctional communication compartments in the Drosophila wing imaginal disk. Dev Biol 102, 130–146. 10.1016/0012-1606(84)90181-7.

55. Weir, M.P., and Lo, C.W. (1982). Gap junctional communication compartments in the Drosophila wing disk. Proc Natl Acad Sci U S A 79, 3232–3235. 10.1073/pnas.79.10.3232.

56. Narciso, C., Wu, Q., Brodskiy, P., Garston, G., Baker, R., Fletcher, A., and Zartman, J. (2015). Patterning of wound-induced intercellular Ca(2+) flashes in a developing epithelium. Phys Biol 12, 056005. 10.1088/1478-3975/12/5/056005.

57. Brodskiy, P.A., Wu, Q., Soundarrajan, D.K., Huizar, F.J., Chen, J., Liang, P., Narciso, C., Levis, M.K., Arredondo-Walsh, N., Chen, D.Z., and Zartman, J.J. (2019). Decoding Calcium Signaling Dynamics during Drosophila Wing Disc Development. Biophys J 116, 725–740. 10.1016/j.bpj.2019.01.007.

58. Balaji, R., Bielmeier, C., Harz, H., Bates, J., Stadler, C., Hildebrand, A., and Classen, A.K. (2017). Calcium spikes, waves and oscillations in a large, patterned epithelial tissue. Sci Rep 7, 42786. 10.1038/srep42786.

59. Warner, A.E., and Lawrence, P.A. (1982). Permeability of gap junctions at the segmental border in insect epidermis. Cell 28, 243–252.

60. Peterson, N.G., Stormo, B.M., Schoenfelder, K.P., King, J.S., Lee, R.R., and Fox, D.T. (2020). Cytoplasmic sharing through apical membrane remodeling. Elife 9. 10.7554/eLife.58107.

61. Monterisi, S., Michl, J., Hulikova, A., Koth, J., Bridges, E.M., Hill, A.E., Abdullayeva, G., Bodmer, W.F., and Swietach, P. (2022). Solute exchange through gap junctions lessens the adverse effects of inactivating mutations in metabolite-handling genes. Elife 11. 10.7554/eLife.78425.

62. Nakai, J., Ohkura, M., and Imoto, K. (2001). A high signal-to-noise Ca(2+) probe composed of a single green fluorescent protein. Nat Biotechnol 19, 137–141. 10.1038/84397.

63. Razzell, W., Evans, I.R., Martin, P., and Wood, W. (2013). Calcium flashes orchestrate the wound inflammatory response through DUOX activation and hydrogen peroxide release. Current biology : CB 23, 424-429. 10.1016/j.cub.2013.01.058.

64. Speder, P., and Brand, A.H. (2014). Gap junction proteins in the blood-brain barrier control nutrient-dependent reactivation of Drosophila neural stem cells. Dev Cell 30, 309–321. 10.1016/j.devcel.2014.05.021.

65. Ho, K.Y.L., Khadilkar, R.J., Carr, R.L., and Tanentzapf, G. (2021). A gap-junction-mediated, calcium-signaling network controls blood progenitor fate decisions in hematopoiesis. Curr Biol 31, 4697–4712 e4696. 10.1016/j.cub.2021.08.027.

66. Ho, K.Y.L., An, K., Carr, R.L., Dvoskin, A.D., Ou, A.Y.J., Vogl, W., and Tanentzapf, G. (2023). Maintenance of hematopoietic stem cell niche homeostasis requires gap junction-mediated calcium signaling. Proc Natl Acad Sci U S A 120, e2303018120. 10.1073/pnas.2303018120.

67. Restrepo, S., and Basler, K. (2016). Drosophila wing imaginal discs respond to mechanical injury via slow InsP3R-mediated intercellular calcium waves. Nat Commun 7, 12450. 10.1038/ncomms12450.

68. Moore, J.L., Bhaskar, D., Gao, F., Matte-Martone, C., Du, S., Lathrop, E., Ganesan, S., Shao, L., Norris, R., Campama Sanz, N., et al. (2023). Cell cycle controls long-range calcium signaling in the regenerating epidermis. J Cell Biol 222. 10.1083/jcb.202302095.

69. Tu, R., Tang, X.A., Xu, R., Ping, Z., Yu, Z., and Xie, T. (2023). Gap junction-transported cAMP from the niche controls stem cell progeny differentiation. Proc Natl Acad Sci U S A 120, e2304168120. 10.1073/pnas.2304168120.

70. Siddiqui, M.A., Rømer, A.M.A., Pinna, L., Ramesh, V., Møller, S.S., Gollavilli, P.N., Turtos, A.M., Varghese, S., Guldborg, L.B., Palani, N.P., et al. (2024). Contact-based thymidylate transfer promotes collective tumor growth. bioRxiv, 2024.2007.2031.606056. 10.1101/2024.07.31.606056.

71. Azpiazu, N., and Morata, G. (2000). Function and regulation of homothorax in the wing imaginal disc of Drosophila. Development 127, 2685–2693. 10.1242/dev.127.12.2685.

72. Furriols, M., and Bray, S. (2001). A model Notch response element detects Suppressor of Hairless-dependent molecular switch. Curr Biol 11, 60–64. S0960-9822(00)00044-0 [pii].

73. Bosveld, F., Bonnet, I., Guirao, B., Tlili, S., Wang, Z., Petitalot, A., Marchand, R., Bardet, P.L., Marcq, P., Graner, F., and Bellaiche, Y. (2012). Mechanical control of morphogenesis by Fat/Dachsous/Four-jointed planar cell polarity pathway. Science 336, 724–727. 10.1126/science.1221071.

74. Bardin, A.J., Perdigoto, C.N., Southall, T.D., Brand, A.H., and Schweisguth, F. (2010). Transcriptional control of stem cell maintenance in the Drosophila intestine. Development 137, 715–724.

75. Loubiere, V., Delest, A., Thomas, A., Bonev, B., Schuettengruber, B., Sati, S., Martinez, A.M., and Cavalli, G. (2016). Coordinate redeployment of PRC1 proteins suppresses tumor formation during Drosophila development. Nat Genet 48, 1436–1442. 10.1038/ng.3671.

76. Ostrowski, K., Bauer, R., and Hoch, M. (2008). The Drosophila innexin 7 gap junction protein is required for development of the embryonic nervous system. Cell Commun Adhes 15, 155–167. 10.1080/15419060802013976.

77. Smendziuk, C.M., Messenberg, A., Vogl, A.W., and Tanentzapf, G. (2015). Bi-directional gap junction-mediated soma-germline communication is essential for spermatogenesis. Development 142, 2598–2609. 10.1242/dev.123448.

78. Ammer, G., Vieira, R.M., Fendl, S., and Borst, A. (2022). Anatomical distribution and functional roles of electrical synapses in Drosophila. Curr Biol 32, 2022–2036 e2024. 10.1016/j.cub.2022.03.040.

79. Bohrmann, J., and Zimmermann, J. (2008). Gap junctions in the ovary of Drosophila melanogaster: localization of innexins 1, 2, 3 and 4 and evidence for intercellular communication via innexin-2 containing channels. BMC Dev Biol 8, 111. 10.1186/1471-213X-8-111.

80. Knecht, W., Munch-Petersen, B., and Piskur, J. (2000). Polyclonal antibodies against the ultrafast multisubstrate deoxyribonucleoside kinase from Drosophila melanogaster. Adv Exp Med Biol 486, 263–266. 10.1007/0-306-46843-3_51.

81. Al Zouabi, L., Stefanutti, M., Roumeliotis, S., Le Meur, G., Boumard, B., Riddiford, N., Rubanova, N., Bohec, M., Gervais, L., Servant, N., and Bardin, A.J. (2023). Molecular underpinnings and environmental drivers of loss of heterozygosity in Drosophila intestinal stem cells. Cell Rep 42, 113485. 10.1016/j.celrep.2023.113485.

82. Schindelin, J., Arganda-Carreras, I., Frise, E., Kaynig, V., Longair, M., Pietzsch, T., Preibisch, S., Rueden, C., Saalfeld, S., Schmid, B., et al. (2012). Fiji: an open-source platform for biological-image analysis. Nat Methods *9*, 676-682. 10.1038/nmeth.2019.

83. Stirling, D.R., Swain-Bowden, M.J., Lucas, A.M., Carpenter, A.E., Cimini, B.A., and Goodman, A. (2021). CellProfiler 4: improvements in speed, utility and usability. BMC Bioinformatics 22, 433. 10.1186/s12859-021-04344-9.

84. Blanco-Obregon, D., El Marzkioui, K., Brutscher, F., Kapoor, V., Valzania, L., Andersen, D.S., Colombani, J., Narasimha, S., McCusker, D., Leopold, P., and Boulan, L. (2022). A Dilp8-dependent time window ensures tissue size adjustment in Drosophila. Nat Commun 13, 5629. 10.1038/s41467-022-33387-6.

